# The vacuolar tauopathy-associated mutation D395G confers redox sensitivity to p97/VCP

**DOI:** 10.64898/2026.04.16.718620

**Authors:** Smruti R. Rout, Martin Kulke, Maxim Droemer, Marcel Wendel, Tat Cheung Cheng, Tristan A. Mauck, Sean Asgari, Antonia A. Nemec, Mikhail Shein, Yusuke Sato, Shuya Fukai, Gregor Witte, Robert J. Tomko, Florian Stengel, Martin Zacharias, Anne K. Schuetz, Eri Sakata

**Affiliations:** Institute for Auditory Neuroscience, University Medical Center Göttingen, 37077 Göttingen, Germany; Institute for Neuropathology, University Medical Center Göttingen, 37073 Göttingen, Germany; Multiscale Bioimaging: from Molecular Machines to Networks of Excitable Cells (MBExC), University of Göttingen, 37073 Göttingen, Germany; Center for Functional Protein Assemblies, Technical University of Munich, 85748 Garching, Germany; Department of Physics, Technical University of Munich, 85748 Garching, Germany; Faculty for Chemistry and Pharmacy, Ludwig-Maximilians-Universität München, 81377 Munich, Germany; Department of Biology, University of Konstanz, 78457 Konstanz, Germany; Konstanz Research School of Chemical Biology, University of Konstanz, 78457 Konstanz, Germany; Department of Biomedical Sciences, Florida State University College of Medicine, 1115 W. Call St., Tallahassee, FL 32306, USA; Department of Chemistry and Biotechnology, Graduate School of Engineering, Tottori University, 4-101 Koyama-cho Minami, Tottori-shi, Tottori 680-8552, Japan; Center for Research on Green Sustainable Chemistry, Tottori University, 4-101 Koyama-cho Minami, Tottori-shi, Tottori 680-8552, Japan; Chromosome Engineering Research Center, Tottori University, 86 Nishi-cho, Yonago-shi, Tottori 683-8503, Japan; Department of Chemistry, Graduate School of Science, Kyoto University, Kyoto 606-8502, Japan; Gene Center and Department of Biochemistry, Ludwig-Maximilians-Universität München, 81377 Munich, Germany

## Abstract

The multifunctional AAA+ ATPase p97/VCP is a pivotal regulator of cellular proteostasis, extracting polyubiquitinated substrates from protein complexes or organelle membranes for proteasomal degradation. Mutations in p97 are linked to a broad spectrum of neurodegenerative disorders, including multisystem proteinopathy and amyotrophic lateral sclerosis. Here, we provide insights into the basis of dysfunction in p97^D395G^, implicated in vacuolar tauopathy, using an integrated structural approach. The mutation destabilizes the interaction network in the transient ADP.P_i_ state previously identified in p97^WT^ and alters the dynamics of the linker between the tandem ATPase domains, resulting in decreased ATPase activity. We further demonstrate that the D395G mutation sensitizes p97 to oxidative stress by enhancing C522 oxidation, thereby perturbing nucleotide binding in D2, and define the functional basis of this oxidative inactivation of p97. These findings reveal redox control as a key regulatory layer of the p97 ATPase cycle and provide a mechanistic framework for how oxidative stress contributes to p97-associated neurodegeneration.

## Introduction

Among ATPases Associated with diverse cellular Activities (AAA+) proteins, p97 is distinguished by its extensive network of cofactors, which couple its unfolding activity to a wide range of intracellular pathways, including protein homeostasis, membrane remodeling, and chromatin regulation ^1,2^. Pathogenic mutations in p97 have been linked to multiple neurodegenerative diseases, i.e., inclusion body myopathy associated with Paget disease of bone and frontotemporal dementia, also known as multisystem proteinopathy (MSP), amyotrophic lateral sclerosis, and vacuolar tauopathy ^3–5^. The best-studied p97 mutations associated with MSP-1 are gain-of-function variants, exhibiting hyperactive ATPase activity, enhanced substrate unfolding, and accumulation of TDP-43 aggregates ^6–8^. In contrast, the recently identified D395G (aspartate to glycine) mutation represents a loss-of-function variant that impairs p97 ATPase activity and is associated with tau deposition in the brain ^9,10^. p97 was identified as a critical tau disaggregase that promotes the clearance of pathological tau aggregates, linking its dysfunction to neurodegenerative mechanisms ^11^.

p97 comprises an N-terminal domain (NTD), two tandem ATPase domains (D1 and D2) connected by a flexible D1-D2 linker, and a short unstructured C-terminal domain (CTD) ^1^. It assembles into a homo-hexameric ring with the D1 ring stacked upon the D2 ring. The NTDs undergo large-scale rearrangements: in the presence of ATP or non-hydrolysable analogs (e.g. ATPγS), they adopt an ‘up’ conformation, whereas following ATP hydrolysis, they transition to a position coplanar with the D1 ring (‘down’ state) ^12–14^. Recently, we reported a cryo-electron microscopy (cryo-EM) structure of p97 in ADP.P_i_ state, representing a reaction intermediate in which neither cleaved phosphate nor ADP has yet been released from the D1 domain. In this state, p97 adopts an NTD-‘down’ conformation, demonstrating that alterations in the interaction network upon ATP hydrolysis stabilize the NTD in ‘down’ position ^15^. Efficient ATP hydrolysis requires coordination between the two ATPase domains, which is mediated in part via the D1-D2 linker ^14,16,17^. Upon substrate engagement, p97 breaks the symmetry of the hexamer and adopts a translocation-active conformation with a spiral-staircase arrangement ^18^. ATP hydrolysis drives conformational changes coupled to pore loop movements, which modulate substrate interactions and facilitate protein unfolding ^18^.

The functional versatility of p97 stems from its dynamic association with numerous co-factors that modulate its activity ^19^, oligomerization state ^20^, and subcellular localization ^21,22^. In addition, a plethora of post-translational modifications including phosphorylation, ubiquitination, and oxidation ^2,23–26^, have been reported to fine-tune p97 activity. Oxidative modifications predominantly target nucleophilic residues such as cysteine, methionine, and tryptophan, with sulfur-based chemistry being most prominent: cysteine thiols readily form disulfide bonds, whereas higher oxidation states yield reversible sulfenylation or irreversible sulfinylation and sulfonylation ^27^. Oxidation of p97 and its homologs Cdc48 in different organisms impairs its activity ^25,26,28^. However, the structural consequences of such oxidative modification remain poorly understood.

In this study, we used cryo-EM to solve multiple structural states of p97^D395G^ in the presence of distinct nucleotides, revealing the intra- and inter-subunits communication network that enables allosteric regulation and providing structural insight into the molecular dysfunction of the p97^D395G^ mutant. We further investigate the impact of oxidative stress on p97 and reveal the mechanistic basis of its inactivation through oxidative modification of a conserved cysteine within the D2 active site, providing further insights into redox-mediated regulation of p97 function under stress conditions.

## Results

### Molecular defects of p97^D395G^ are mechanistically distinct from MSP mutants

D395 is located in the small subdomain of the D1 ATPase, capping an α helix_397-402_ (Fig. 1a and Extended Fig. 1a). Both the D1 and D2 nucleotide binding pockets are distant from D395 (Extended Fig. 1a). The pathogenic mutation D395G reduces ATPase activity ^9^, which is coupled with decreased unfolding activity toward a model substrate, polyubiquitinated Ub_4_-mEOS3.2 (Fig. 1b). To explore the structural impact of the mutation, we first employed methyl-detected NMR spectroscopy using the ND1L (p97_1-480_) hexamer, comprising of NTD, D1, and D1-D2 linker ^29^. Wild type p97 (p97^WT^) ND1L adopts an NTD-‘down’ conformation coplanar with the D1 ring in the presence of ADP, and an ‘up’ conformation in the presence of ATP analogues ^29^. Either conformation can be readily identified from the NMR signal positions of reporter residues located in the NTD. The ADP-bound forms of canonical MSP-1 mutants like p97^R95G^ exhibit fast exchange (> 2000 /sec) between ‘up’ and ‘down’ states, resulting in averaged NMR signals (Fig. 1c). In contrast, p97^D395G^ displays NMR spectra largely identical to the corresponding p97^WT^ state regardless of the investigated nucleotide state; ADP, and ATPγS-bound states as well as in the presence of an ATP regeneration system (ATP^reg^). Interestingly, the subset of residues that report on NTD position (V87, V116, V154, V161, and V176) displays a doubling of methyl correlations corresponding to both the ‘up’ (minor signal) and ‘down’ (major signal) conformations in ADP-bound form (Fig. 1c). This indicates the presence of a slow exchange regime between the two conformations with the ‘up’ state populated to a much lower extent, which is distinct from the observation in MSP-1 mutants ^30^. Importantly, p97^D395G^ retains an intact ‘up’ conformation in the presence of ATPγS, confirming structural responsiveness to ATP analogue binding (Fig. 1c). An in-depth analysis of the spectrum in the ADP state revealed that only methyl probes near the D395G mutation site experience significant chemical shift perturbations, indicating that the D395G mutation induces no major structural rearrangement in the D1 domain (Extended Fig. 1d).

**Figure 1:**
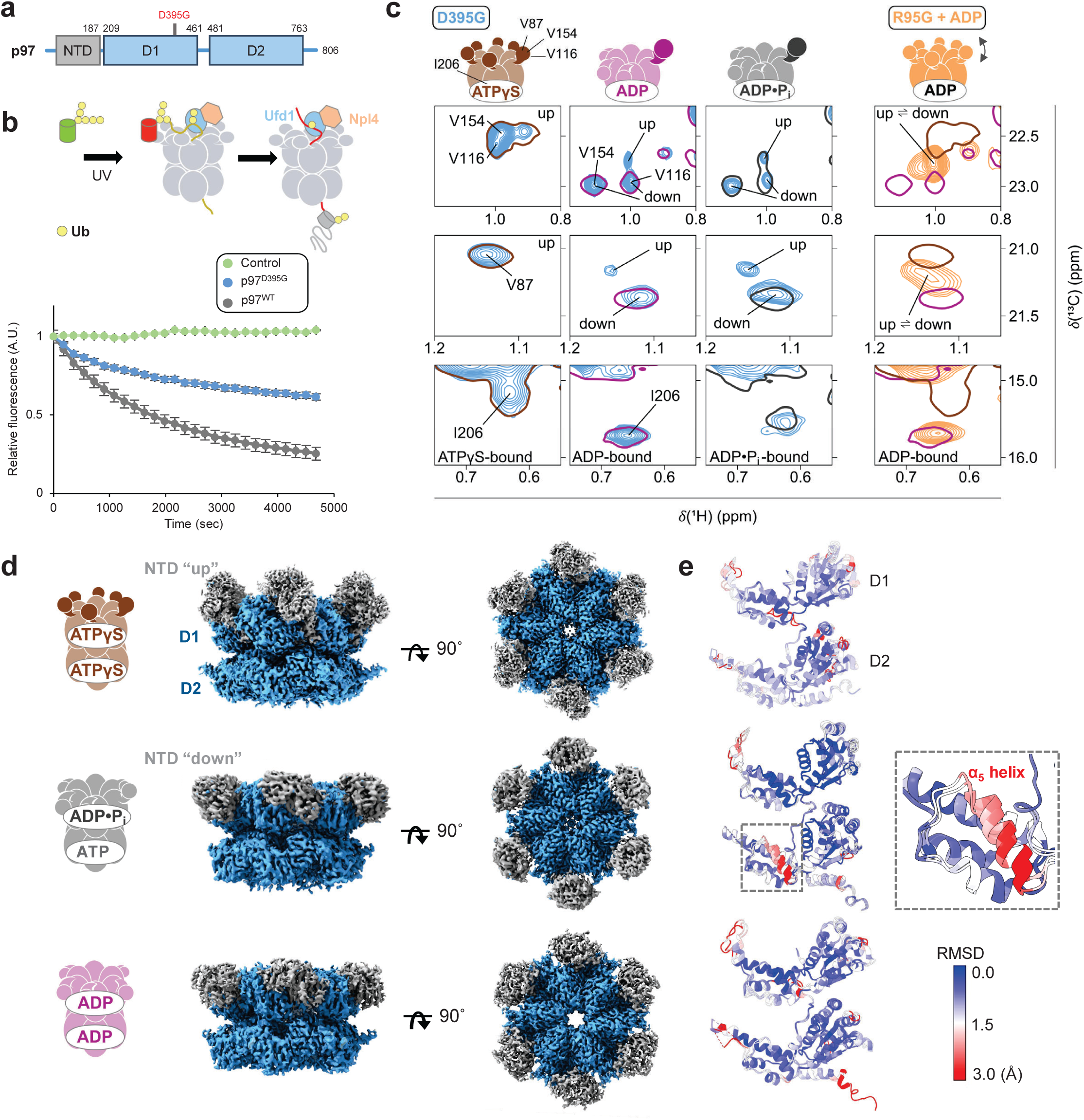
p97^D395G^ mutant exhibits subtle conformational alterations. **a**, Domain structure of p97, highlighting the D395G mutation located in the D1 domain. **b,** Schematic representation of preparation of ubiquitinated Ub□-mEOS3.2 for the p97 substrate unfolding assay (top). Green Ub□-mEOS3.2 was polyubiquitinated using E1, E2-25K, and ubiquitin, and subsequently photoconverted to its red form under UV illumination, as previously described ^8^. The purified substrate was then used in unfolding assays in the presence of the Ufd1-Npl4 complex. Compared with p97^WT^ (grey), p97^D395G^ (blue) exhibits reduced substrate unfolding activity (bottom). All measurements were performed with at least three technical replicates. **c,** NMR characterization of p97^D395G^ using the ND1L construct, comprising the NTD, D1 domain and linker. Key probes for NTD conformation and bound nucleotide are indicated in the schematic above. ATPγS induces an NTD ‘up’ conformation, similar to p97^WT^. In the presence of ADP and the ATP^reg^, predominantly ‘down’ with a minor ‘up’ conformation is observed, in contrast to p97^WT^. For comparison, spectra of the prototypical MSP-1-associated mutant p97^R95G^ in presence of ADP are shown, which display signals at an averaged position between NTD ‘up’ and ‘down’ conformation, indicating fast conformational exchange on the NMR time scale. This establishes that p97^D395G^ displays distinct molecular defects from MSP-1-type mutants. **d,** Cryo-EM reconstructions of p97^D395G^ under different conditions (ATPγS, ATP^reg^, and ADP). The N-terminal domain (NTD) is colored grey, and the D1 and D2 ATPase domains are shown in blue. Schematic representations on the left indicate the corresponding nucleotide-bound states. **e,** Backbone RMSD of a p97^D395G^ protomer relative to p97^WT^ in different nucleotide states, calculated without the NTD. p97^D395G^ structures in ATPγS, ADP, and ADP.P_i_ states were aligned using the main chain atoms of the D1 and D2 domains to the published p97^WT^ models in the ATPγS (PDB: 5FTN), ADP (PDB: 5FTK), and ADP.P_i_ (PDB: 8OOI) states, respectively. The inset shows a magnified view of the α_5_ helix in the D2 domain of p97^D395G^ in ADP.P_i_ state.

Molecular dynamics simulations (MDS) of full-length p97 in the ATP-bound state showed that the D395G mutation induces only minor backbone root mean square deviation (RMSD) changes in the ATPase core, except in the C-terminal region of the D2 domain, which displays high conformational flexibility (Extended Fig. 1e-f, Supplementary video 1). Correlation map analysis identified allosteric interactions within structured regions of the NTD, D1, and D2 domains, as well as between neighboring monomers (Extended Fig. 1g). Together, these results suggest that p97^D395G^ alters the conformational landscape of p97 only in a subtle manner.

Next, we solved the cryo-EM structures of p97^D395G^ in the presence of three different nucleotides: ADP, ATPγS, and ATP in ATP^reg^ system (Fig. 1d, Extended Fig. 2-3). Similar to p97^WT^, p97^D395G^ adopts an all-‘down’ NTD conformation in the presence of ADP (Extended Fig. 3a). In contrast, p97^D395G^ NTD shows structural heterogeneity both in the ATPγS and ATP^reg^ conditions. In ATPγS, only 22% of particles exhibit the all-‘up’ NTD conformation (Extended Fig. 3b), whereas p97^WT^ particles are exclusively in NTD ‘up’ state. In ATP^reg^ condition, ∼80% particles adopt the all-‘down’ conformation, while the remaining particles show flexible NTDs in two of six subunits (Extended Fig. 2a). Thus, our data indicate that the p97^D395G^ mutant exhibits defects in nucleotide-dependent inter-domain motions of the NTD. For further comparison with p97^WT^ structures, we refined p97^D395G^ classes corresponding to all-‘down’ state in ADP and ATP^reg^ and all-‘up’ state in ATPγS to build 3D models. Both in the ADP and ATPγS states (Fig 1E), only peripheral regions and flexible loops show structural diversity but the no difference is observed in the nucleotide-binding pocket (Extended Fig. 4). Under ATP^reg^ conditions, we observed a post-hydrolysis ADP.P_i_ state in the D1 pocket, whereas ATP was bound in the D2 pocket (Extended Fig. 4), as previously observed in p97^WT^ ^15^. This configuration (ADP.P_i_ in D1 and ATP in D2) is referred to as ADP.P_i_ state hereafter. Comparison of the backbone RMSD of an isolated p97^D395G^ protomer with respect to p97^WT^ demonstrated that the mutation does not extensively alter the structure (Fig. 1e). Interestingly, a downward displacement of helix α_649-662_ (referred to as α_5_ helix hereafter) was seen in the large subdomain of D2 in the ADP.P_i_ state (Fig. 1e, inset). This allosteric and nucleotide state-specific rearrangement indicates that the mutation primarily perturbs this state, which was therefore the focus of subsequent analysis.

### The D395G mutation affects nucleotide coordination in the D2 domain

The nucleotide-binding pockets of p97^D395G^ both in the ADP and ATPγS-bound states are indistinguishable from p97^WT^ (Extended Fig. 4a). In p97^D395G^ in ADP.P_i_, however, the D2 pocket is majorly affected but not the D1 pocket (Fig. 2a-b). Walker B E578 and arginine finger R635 in the D2 pocket of p97^D395G^ in ADP.P_i_ adopt two distinct side-chain orientations, which are absent in p97^WT^ (Fig. 2c, Extended Fig. 5a). Interestingly, one E578 rotamer is stabilized through interaction with K584, which repositions it away from the catalytic Mg^2+^ ion (Fig. 2c). MDS further supports this, illustrating that the contact between E578 and K584 is significantly stabilized in p97^D395G^ (Fig. 2d). This regulation of the Walker B E578 by its neighboring lysine residue resembles the “glutamate switch” mechanism described in AAA+ ATPases, which transiently suppresses their activities ^31,32^. Besides, the coordination between the ATP adenine ring and conserved S652 at the D2 α_5_ helix in p97^WT^ in ADP.P_i_ is lost in the p97^D395G^, due to a 6.7 Å downward shift of α_5_ helix (Fig. 2b, 2e, Supplementary video 2). Both K584A and S652A mutations decrease ATPase activity, supporting its regulatory role in Mg^2+^ and nucleotide coordination, respectively (Fig. 2f). Surface plasmon resonance analysis showed comparable nucleotide affinities for p97^WT^ and p97^D395G^, emphasizing that the reduced ATPase activity of p97^D395G^ arises from slower catalytic turnover due to suboptimal configuration of nucleotide rather than improper nucleotide binding (Extended Fig. 5c-d).

**Figure 2:**
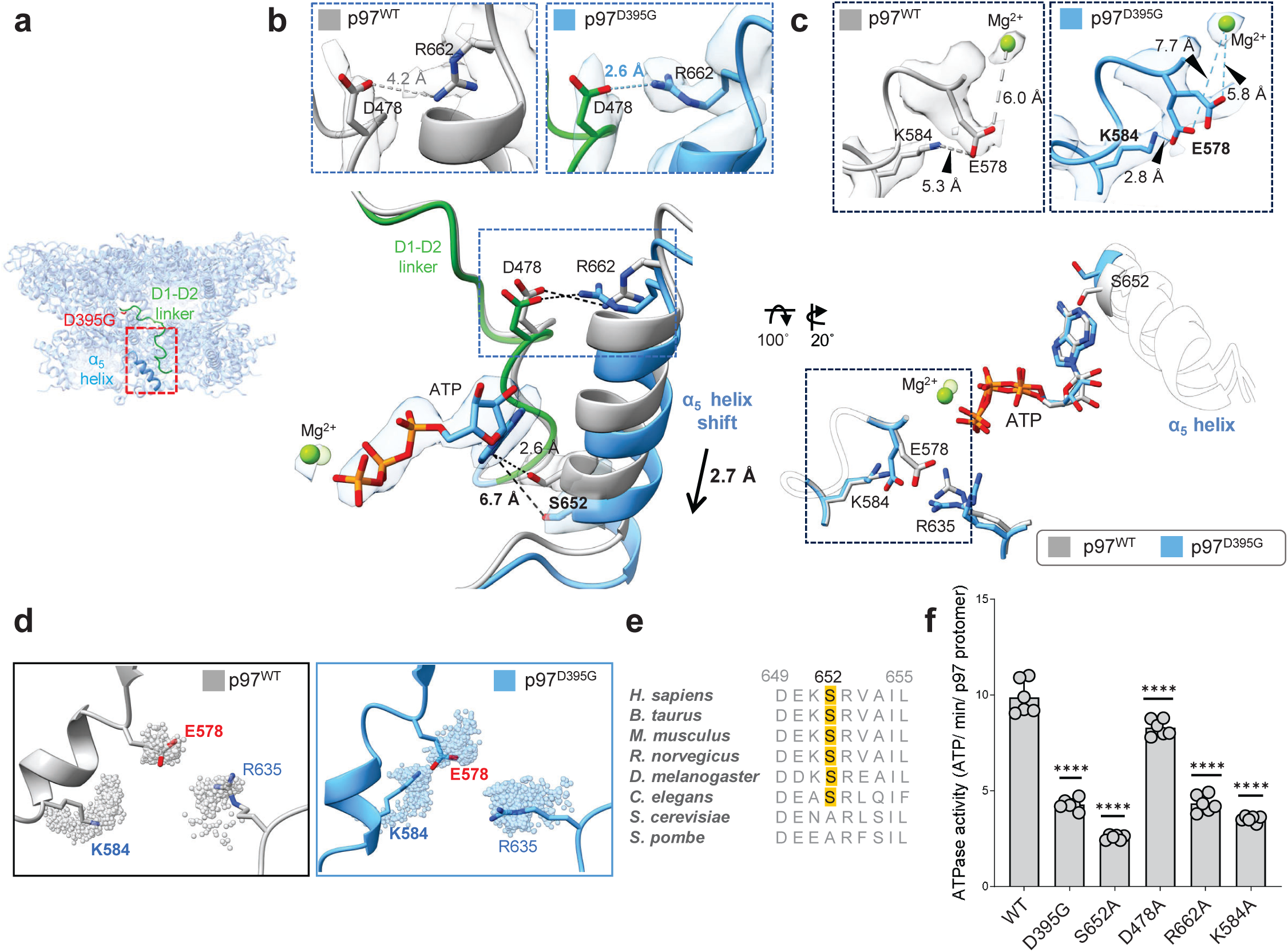
D395G mutation impairs nucleotide coordination in D2 active site in the ADP.P_i_ state. **a**, Structure of p97^D395G^ in the ADP.P_i_ state, highlighting the α_5_ helix of the D2 domain, the D1-D2 linker, and the D395G mutation (colored blue, green, and red, respectively). **b,** The α_5_ helix shifts downward by 2.7 Å in p97^D395G^ relative to p97^WT^ (grey), resulting in improper nucleotide coordination by S652, which is positioned 6.7 Å away in the mutant. This helical displacement is stabilized by an upstream interaction between D478 and R662 in p97^D395G^. Differences in the D478-R662 interaction between p97^WT^ and p97^D395G^ are shown in the zoomed-in view above together with the corresponding cryo-EM density map. **c,** In p97^D395G^, Walker B residue E578 adopts two rotameric configurations not seen in p97^WT^. The stabilization of E578 by K584 results in E578 facing away from the Mg^2+^ atom. Structural models with corresponding cryo-EM density map are shown above. **d,** Residue outermost atom positions of K584, E578, and R635 are shown as spheres from 1000 snapshots taken every 1 ns from 1 μs of MDS, illustrating their conformational flexibility. The contact between E578 and K584 is significantly increased in p97^D395G^ compared to p97_WT_. **e,** S652 is highly conserved in metazoans, with alanine at the equivalent position in yeast. **f,** ATPase activity assays were performed on p97 mutants, targeting residues altered by the D395G mutation. K584A reduces ATPase activity to levels comparable to p97^D395G^, highlighting its role in regulating the D2 active site. Similarly, D478A and R662A likely disrupt the active-site configuration, resulting in reduced ATPase activity. All measurements were performed with at least three technical replicates (Asterisks indicate statistical significance calculated using one-way ANOVA: P < 0.0001).

### D1-D2 linker dynamics in inter-protomer communication in p97^D395G^

The displaced α_5_ helix in p97^D395G^ in ADP.P_i_ is stabilized by an electrostatic interaction between R662 and residue D478 located in the D1-D2 linker (Fig. 2b, inset). Structural changes observed in the D2 active site imply that the D395G mutation in D1 acts allosterically by altering D1-D2 linker dynamics (Extended Fig. 6a). MDS shows unchanged secondary structure near the mutation site but subtle local rearrangements. In MDS, the D395G substitution shifts the adjacent residues V394 and L396 closer together by ∼1 Å (Fig. 3a-b), altering the interaction network of the hydrophobic cluster 1 (D393, T448, M449, V394). This rearrangement displaces the α_9_ helix (D1_449-458_) away from the loop containing D395, thereby shifting the hydrophobic cluster 2 (L464, S463, H404, V407, H406). Consequently, the D1-D2 linker drifts further away from the loop_404-407_, increasing the conformational flexibility of R465 (Fig. 3a) and modifying its interactions with D607 and D564 in D2. Overall, all movements originated from the D395G substitution create extra space for the linker movement (Fig. 3a).

**Figure 3:**
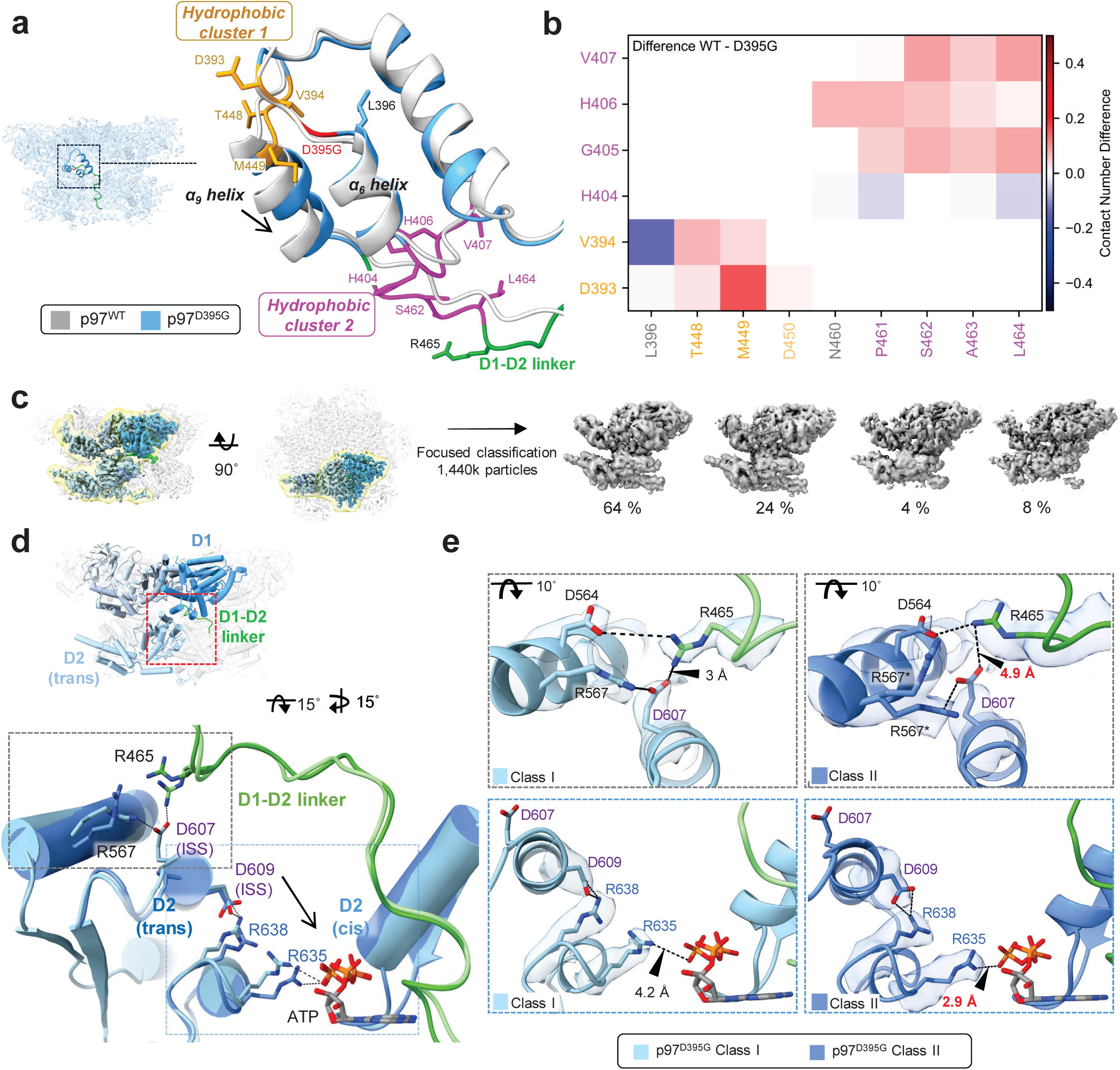
Allosteric communication via D1-D2 linker in p97^D395G^. **a,** Superposition of the region surrounding D395G (red) derived from MDS. Models of p97^WT^ and p97^D395G^ are shown in grey and blue with the D1-D2 linker highlighted in green. The two hydrophobic cluster that undergo pronounced rearrangements upon mutation are shown in gold and pink, with their constituent residues. **b,** Contact differences between p97^WT^ and p97^D395G^ for the relevant residues in the region around D395G including residues from the two hydrophobic clusters are colored in gold and pink. Red and blue colors indicate more pronounced contacts in p97^WT^ and p97^D395G^, respectively. **c,** Schematic representation of focused classification using a mask (yellow) applied to the interface between D1 and the adjacent D2 subunit (trans). Two major classes (Class I, 64 % and Class II, 24 %) were identified in p97^D395G^. **d,** Superposition of two classes obtained from the focused classification of p97^D395G^. The two classes differ markedly in the orientation of R465 within the D1-D2 linker (green). Class I is shown in pale blue with its D1-D2 linker in pale green, and Class II in dark blue with its D1-D2 linker in dark green. Inter-protomer communication proceeds via D1-D2 linker to ISS motif, which regulates positioning of arginine finger residues (arrow). **e,** Zoomed-in views of interactions at the D1-D2 interface. Models are displayed as ribbon representation with the corresponding cryo-EM density. R465 interacts with ISS motif residue D607 (grey dashed box), while ISS residue D609 interacts with arginine finger R638 (blue dashed box). D537 in p97^D395G^ shows rotamer (*) conformation in class II. Models are displayed as ribbon representation with the corresponding cryo-EM density. In Class I, stronger R465-D607 interaction correlate with weaker engagement of R635 with nucleotide whereas in Class II, weakened R465-D607 (red label) disrupts the D609-R638 interaction, consequently enhancing R635 interaction with nucleotide (red label).

Focused classification on C6-symmetry-expanded particles of both the p97^D395G^ and p97^WT^ datasets in ADP.P_i_ revealed distinct linker dynamics. p97^WT^ in ADP.P_i_ particles yielded structurally similar classes (Extended Fig. 6c, d), whereas p97^D395G^ in ADP.P_i_ particles were separated into two major classes (class I and class II, ∼64% and ∼24%, respectively) (Fig. 3c, Supplementary video 3). These two classes are distinguished by the orientation of R465 in the D1-D2 linker towards the conserved inter-subunit signaling (ISS) motif in the adjacent D2 subunit (Fig. 3d). The ISS coordinates inter-protomer ATP hydrolysis ^2^. In class I, R465 interacts with ISS D607 in trans, stabilizing D609-R638 coordination but weakening Arg finger R635 interaction with ATP (Fig. 3e). In class II, R465 instead contacts D564 of the trans protomer and no longer coordinates D607 (Fig. 3e). This rearrangement, propagated via the ISS, disrupts D609-R638 coupling, and consequently enhances nucleotide coordination by R635 (Fig. 3e). Together with MDS, the observed differences in the D2 arrangement suggest that the D395G mutation alters the conformational equilibrium of interface residues in the trans D2 subunit, highlighting an allosteric intra-protomer communication between the D1-D2 linker and trans Arg finger R635, mediated by the trans-acting R465.

### D395G mutation enhances sensitivity to oxidation

In the course of purification, we observed a gradual decline in ATPase activity of p97^D395G^. Given that p97 was reported to undergo inactivation upon irreversible oxidation ^25,26^, it prompted us to further investigate how oxidative modification alters its activity and structure. For *in-vitro* oxidation, hydrogen peroxide (H_2_O_2_) was used as an oxidant ^33^ and the oxidized p97 proteins were purified by size exclusion chromatography (Extended Fig. 7b). Treatment with 25 µM H_2_O_2_ reduced ATPase activity of p97^WT^ to ∼45% of untreated levels, whereas p97^D395G^ retained only ∼25% (Fig. 4a). The ATPase activities of p97^WT^ and p97^D395G^ recovered to ∼85% and ∼57%, respectively, indicating the presence of irreversible oxidation (Extended Fig. 7c). Similar decline in ATPase activity was observed with oxidant diamide (Extended Fig. 7d). In contrast, the MSP-1 p97^R95G^ mutant at the NTD–D1 interface maintained ∼ 60% activity, indicating that the hyperactivated MSP-1 mutant is more resistant to oxidation (Extended Fig. 7e-f). These results suggest that the p97^D395G^ mutation increases sensitivity to oxidation, extending previous observations of enhanced sensitivity to salt and thermal stress ^9^.

**Figure 4:**
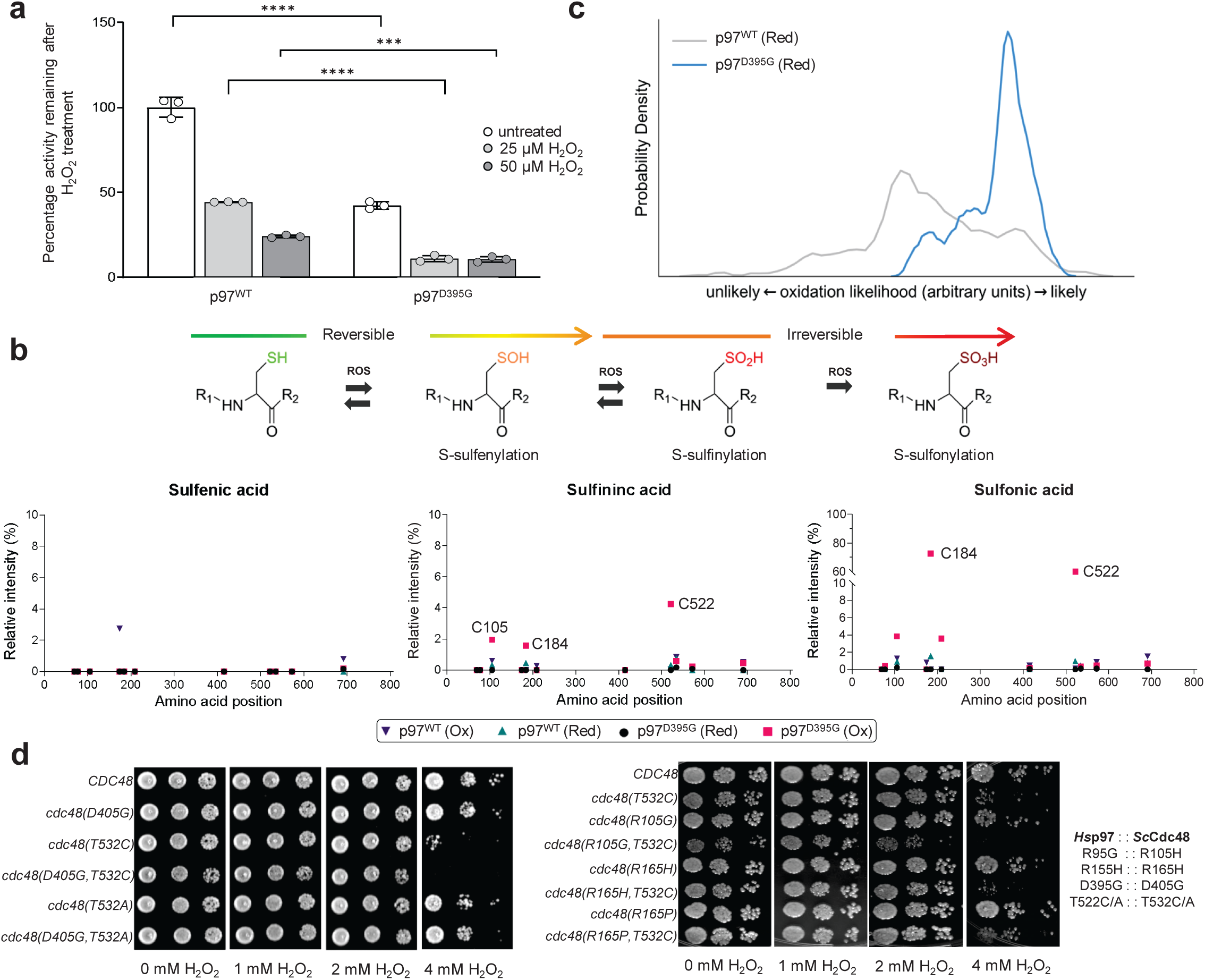
D395G mutation enhances sensitivity to oxidation. **a**, ATPase activity assays after oxidant H_2_O_2_ treatment shows that p97^D395G^ is more susceptible to oxidation. In presence of 25 μM H_2_O_2,_ p97^WT^ retains 45% of activity while p97^D395G^ retains only 25% activity. All measurements include at least three technical replicates (Statistical significance was calculated using two-way ANOVA with factors mutation and oxidation treatment. Asterisks indicate significance levels (P < 0.001(***), P < 0.0001 (****)). b, Mass spectrometry on protein samples oxidized with 25 μM H_2_O_2_ reveals that p97^D395G^ accumulates relatively higher levels of irreversible oxidations – sulfinic acid (C105, C184 and C522) and sulfonic acid (C522 and C184). c, Conformational ensembles of p97^WT^ (grey) and p97^D395G^ (blue) mapped to the likelihood of C522 oxidation to sulfonic acid. The likelihood is represented by a collective variable derived from atomic coordinates surrounding C522 and optimized to distinguish free oxidation energy differences of distinct conformational states. Further details are provided in the methods section. (d) Combining pathogenic mutations of Cdc48 (yeast p97 homolog) with a Cys residue at the D2 nucleotide-binding pocket (T532C substitution) confers oxidation sensitivity in yeast. Growth assays were performed at 30°C for two days with the indicated H_2_O_2_ concentration.

To identify site-specific modifications, oxidized p97 proteins were trypsin-digested, and the modifications of the peptides were analyzed by mass spectrometry (MS). Under identical conditions, p97^D395G^ exhibited higher levels of cysteine oxidation than p97^WT^, with notably increased irreversible modifications (sulfinic and sulfonic acids) at C105, C184, C209, and C522 (Fig. 4b). In the p97^D395G^ mutant, the conserved C522 within the D2 active site shows a high level of oxidation, consistent with earlier cellular assays demonstrating that C522 is highly responsive to oxidative stimuli (Noguchi et al., 2005; Astier et al., 2012). In addition, C184 located in the NTD at the NTD-D1 interface undergoes substantial sulfonic acid modification. Our MS data highlight the D395G mutation indeed enhances oxidative susceptibility (Fig. 4b). Additionally, serine substitution at C184 and C522 residues reduces ATPase activity, indicating their roles in structural stabilization (Extended Fig. 7e). Notably, the oxidation-induced decline in ATPase activity was attenuated in the C522S mutant, whereas it was preserved in the C184S mutant (Extended Fig. 7f). Our findings indicate that oxidation of C522 plays a more critical role in the loss of ATPase activity under oxidative conditions.

Free energies for the C522 oxidation were calculated from MD trajectories using thermodynamic integration (TI) on representative structures of p97^WT^ and p97^D395G^.To enable the estimation of oxidation free energies across all trajectory frames, including those not explicitly calculated by TI, a collective variable, was constructed that describes the oxidation free energy based on the atomic positions of C_α_ atoms near the active center (Extended Fig. 9d), Projecting the trajectories along this collective variable reveals that the p97^D395G^ variant more frequently sampled states with higher oxidation likelihood than p97^WT^ (Fig. 4c), indicating increased susceptibility of the mutant to oxidative modification.

In yeast Cdc48, cysteine 522 of human p97 is replaced by threonine (Extended Fig. 9c). We substituted *S. cerevisiae* T532 (analogous to p97 C522) with cysteine and examined the oxidative stress sensitivity. The D405G substitution (analogous to p97 D395G) alone did not confer oxidative sensitivity, but the double mutant Cdc48^D405G^, ^T532C^ displayed increased sensitivity, supporting that D395G mutation enhances the oxidation sensitivity of C522 (Fig. 4d). MSP-1 mutants combined with T532C mutation showed variable responses; only R105G mutant (analogous to p97 R95G) exhibited a growth defect under oxidative stress (Fig. 4d). However, its ATPase activity was not largely impaired by oxidation (Extended Fig. 7e-f), indicating differential vulnerability to oxidative stress. The growth defect of Cdc48^T532C^ at elevated temperatures was further exacerbated by mutations in Cdc48 cofactors such as Ufd1, Vms1, and Otu1 ^34–36^, suggesting that gain-of-function mutant Cdc48^T352C^ adds an auxiliary function to the Cdc48-dependent protein degradation pathway in yeast (Extended Fig. 7g).

### Heterogenous states observed upon oxidation in p97^D395G^

To address how oxidative modifications remodel the nucleotide-binding sites and alter its catalytic activity, structural analysis of oxidized p97^WT^ and p97^D395G^ was performed in the ATP^reg^ buffer. Oxidized p97^D395G^ populated multiple conformational classes with heterogeneous NTD positioning (Fig. 5a), yielding six distinct conformations with zero to five protomers in the ‘up’ conformation. In contrast, oxidized p97^WT^ remained restricted to the NTD ‘down’ state (Fig. 5a-b). NMR analysis of the p97^D395G^ ND1L construct showed that, in the presence of ADP, H_2_O_2_ exposure progressively shifted the NTD ‘down’ to the ‘up’ state, evidenced by splitting of the signals for Val residues 87, 116, 154, and 133, indicating slow or no exchange between the conformations (Fig. 5c). Because p97^D395G^ ND1L lacks the D2 domain containing C522, the observed shift to the NTD ‘up’ state is likely driven by oxidative modification within the NTD or D1 domains. Given that the highly abundant oxidation site at C184 is located at the NTD-D1 interface, its oxidation may underlie the conformational transition to the ‘up’ state. Hereafter, “oxidized” and “reduced” are abbreviated as “Ox” and “Red,” respectively.

**Figure 5:**
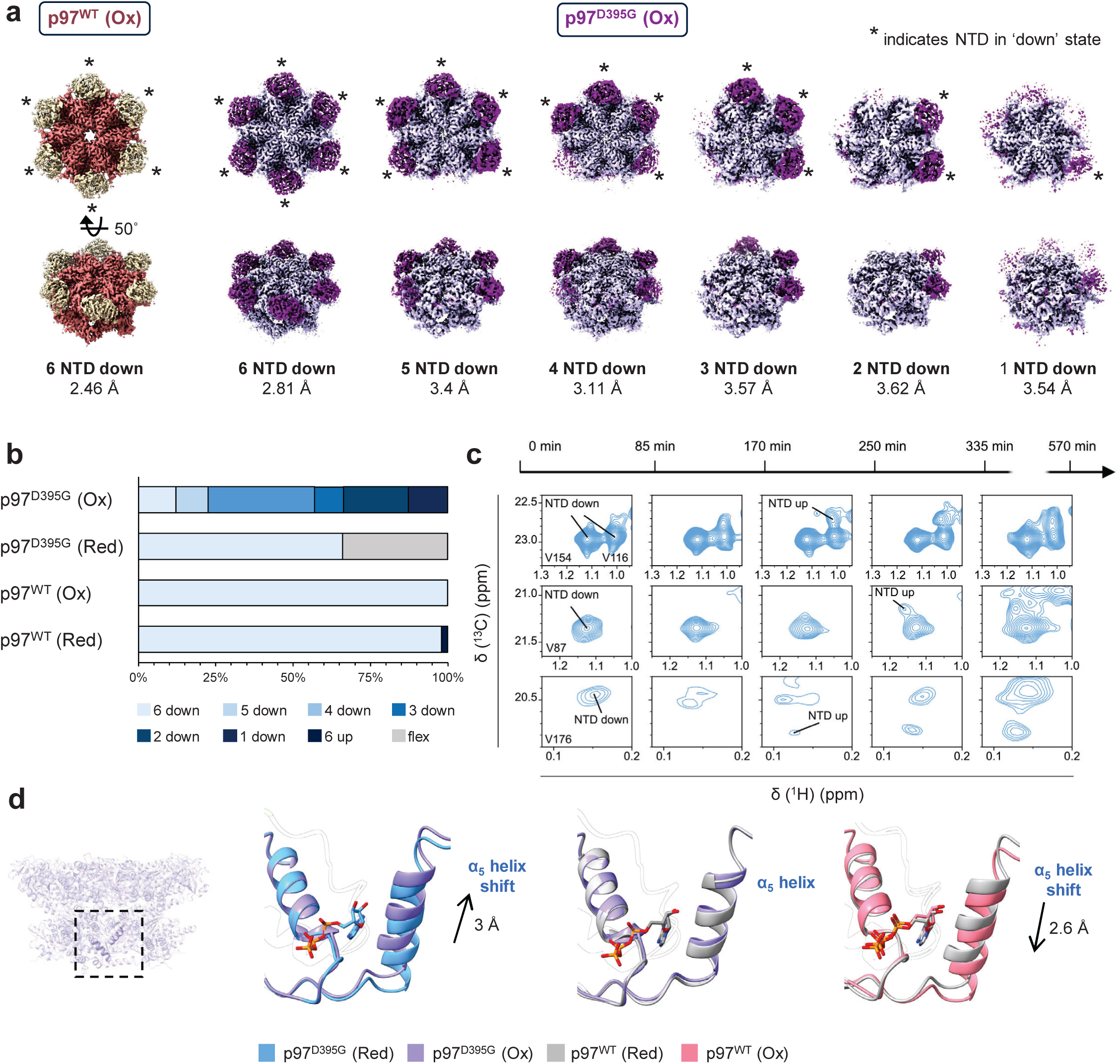
Heterogenous classes observed upon oxidation of p97^D395G^. **a,** Under ATP^reg^ conditions, p97^D395G^ (Ox) is distributed into six distinct cryo-EM classes, reflecting substantial heterogeneity in NTD (purple) positioning. Strikingly, p97^WT^ (Ox) remains conformationally uniform, populating a single NTD (light yellow) ‘down’ class. All reconstructed classes are shown here together with the corresponding map resolutions. The D1-D2 rings of p97^WT^ and p97^D395G^ are colored salmon and light purple, respectively. **b,** Population distribution of p97 conformations based on NTD positioning determined by cryo-EM. ‘Flex’ denotes a class in which one NTD adopts an intermediate, flexible position between the ‘up’ and ‘down’ states. **c,** Real-time NMR methyl correlation spectra recorded on 100 μM p97^D395G^ ND1L protein in the presence of ADP at different time points after addition of 12 mM H_2_O_2_. Gradual splitting of signals reporting on the NTD conformation indicates a transition from all-NTD ‘down’ to mixed ‘up’ and ‘down’ conformations as the oxidation progresses. **d,** Superimposition of p97^D395G^ (Ox) with p97^WT^ (Red) shows recovery of the wild-type-like α_5_ helix conformation. In contrast, comparison of p97^WT^ (Ox) reveals a pronounced displacement of the α_5_ helix in the opposite direction within the D2 ATPase domain. The PDB model 8OOI was used for p97^WT^ (Red).

Among the six classes, the all-NTD ‘down’ class showed the highest resolution in both p97^WT^ (Ox) and p97^D395G^ (Ox), so we further refined these to compare the two structures. Comparison of the oxidized and reduced forms of p97^D395G^ revealed an upward shift of the α_5_ helix in the D2 domain by ∼3 Å (Fig. 5d). This upward shift regains the α_5_ helix conformation of the reduced p97^WT^. Conversely, p97^WT^ (Ox) showed a downward shift of the α_5_ helix by ∼2.6 Å (Fig. 5d). Because the positioning of the α_5_ helix governs nucleotide coordination by S652 and regulates D2 catalysis, proper placement of the α_5_ helix is likely a key determinant of D2 ATPase activity.

### Oxidation-induced conformational rearrangement of C522 prevents nucleotide binding at the D2 active site

Strikingly, the D2 active site pocket of oxidized p97^D395G^ lacked any EM density corresponding to nucleotide, whilst the D1 pocket exhibited the ADP.P_i_ state (Fig. 6a, Extended Fig. 9b). This absence was consistent in all six D2 active sites across every p97^D395G^(Ox) class, irrespective of NTD conformation, suggesting that oxidation of C522 underlies the defective ATPase activity. Strikingly, structural comparison of the D2 active sites between p97^D395G^(Ox) and p97^D395G^ (Red) revealed a pronounced reorientation of C522 (Fig. 6b).

**Figure 6:**
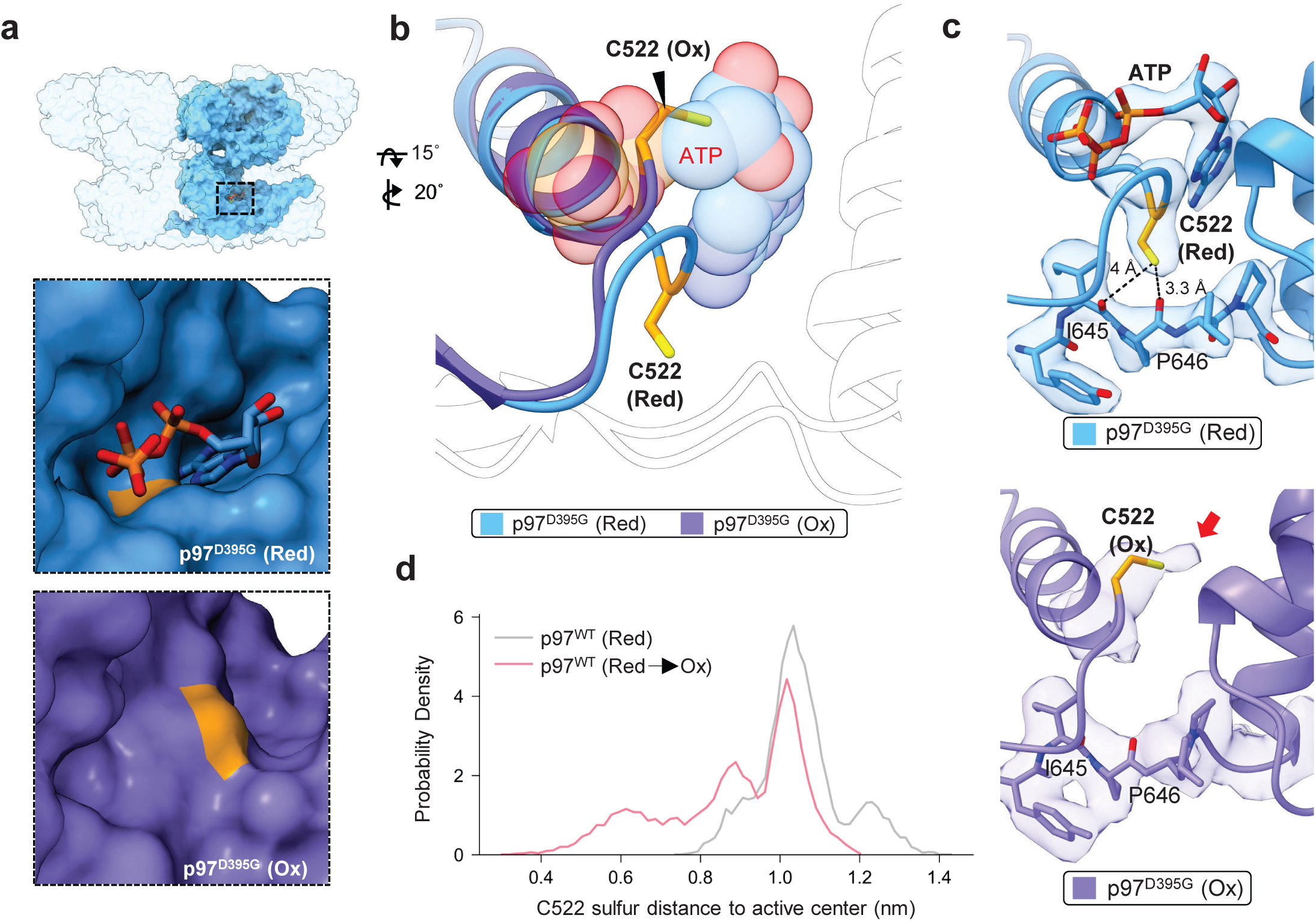
Oxidation-induced conformational rearrangement of C522 prevents nucleotide binding in p97^D395G^ (Ox). **a**, Surface representation of the D2 active site pocket showing absence of bound nucleotide in p97^D395G^ (Ox). p97^D395G^ (Red) in blue, p97^D395G^ (Ox) in purple and C522 in orange. **b,** Superimposition of oxidized p97^D395G^ (purple) and nucleotide-bound reduced form of p97^D395G^ (blue). The bound nucleotide is rendered as transparent spheres to highlight steric repulsion with oxidized C522 in the D2 active site. **c,** *Top:* C522 in the D2 active site of p97^D395G^ (Red) with corresponding EM density. *Bottom*: In p97^D395G^ (Ox), C522 adopts a flipped conformation driven by electrostatic repulsion from the neighboring backbone carbonyl moieties of I645 and P646. The EM density map reveals additional density protruding from C522, indicative of oxidative modification. **d,** Histograms of distances in MDS between C522-S□ and the D2 ATP binding pocket for p97^WT^ in the ‘up’ state with reduced (grey) and oxidized (pink) C522. The atom definition for the active center can be found in the Methods. For both reduced and oxidized C522, MDS were initiated from the nucleotide-free, reduced p97^WT^ structure. Upon oxidation, the C522 side chain reorients towards the D2 pocket, sterically blocking nucleotide binding.

In the reduced form, the C522 thiol is oriented toward the backbone carbonyl groups of I645 and P646. Upon oxidation, C522 flips away from this negatively-charged patch and reorients into the nucleotide binding pocket (Fig. 6b-c). Consistent with this conformational change, an additional EM density protruding from the C522 sulfur atom is observed (Fig. 6c, bottom), supporting the presence of an oxidative modification on the thiol moiety that likely introduces a negatively charged sulfinic or sulfonic group, as identified by MS (Fig. 4c). This modification likely generates electrostatic repulsion with the I645-P636 backbone carbonyls and thereby repositions C522 to occlude the nucleotide binding (Supplementary video 4). Subsequently, C522 in a randomly selected structure from the reduced conformational ensemble of p97^WT^ was converted to the oxidized form (sulfonic acid), and the side chain reorientation of the C522 residue was tracked by MDS. Oxidation causes C522 to reorient toward and obstruct the nucleotide-binding pocket (Fig. 6d), in agreement with our cryo-EM data. C522 is conserved in metazoans but replaced by threonine in yeast, suggesting evolutionary acquisition of a cysteine-based redox sensor that enables higher eukaryotes to modulate p97 D2 activity in response to oxidative stress.

## Discussion

As a central hub in cellular protein homeostasis, p97/Cdc48 coordinates cellular functions related to protein degradation ^37–39^, membrane dynamics ^40^, and chromatin regulation ^41^. Pathogenic mutations in p97 are linked to a spectrum of progressive neurodegenerative disorders ^4,5^, yet how these mutations perturb function remains not fully understood. Previous studies have shown that p97 MSP-1 mutants exhibit different structural dynamics, suggesting that dysregulated conformational behavior of p97 may underlie disease ^8,30^. In this study, we investigated the structural basis of reduced ATPase activity and conformational dynamics in the p97^D395G^ mutant, a variant associated with late-onset vacuolar tauopathy ^9,10^. Additionally, we showed that oxidative modification of p97 introduces an additional layer of regulation that directly impacts nucleotide coordination and conformational dynamics.

Our cryo-EM analysis of p97^D395G^ reveals structural alterations in the NTD switch and in nucleotide coordination within the D2 domain. Although no prominent structural alteration in the presence of ADP (Fig. 1d), distinct coordination of the NTD positions was observed both under the ATPγS and ATP^reg^ conditions, indicating that the D395G mutation affects coupled NTD-conformational shits upon ATP hydrolysis in D1. Furthermore, structural comparison between p97^WT^ and p97^D395G^ shows subtle differences in both ATPγS and ADP, whereas under ATP^reg^ condition, the mutation induces a displacement of the D2 α_5_ helix. This shift perturbs the D2 active site despite the mutation residing in D1, enforcing a suboptimal configuration in which improper nucleotide coordination by S652 contributes to reduced ATPase activity. Notably, S652 localized in the α_5_ helix, is a proposed stress-responsive phosphorylation site ^23^, suggesting that D2 activity may be subject to regulatory tuning through post-translational modification of this structural element. Such a regulatory mechanism would align with the dominant contribution of D2 to the overall ATPase activity of p97 ^42^. Consistent with this, the vacuolization phenotype observed in D2 Walker A mutation (K524A) ^43^ closely resembles neuronal vacuolization phenotype in p97^D395G^ patient tissue ^9^, reinforcing the correlation between D395G mutation and D2 domain catalytic dysfunction.

The reduced ATPase activity of p97^D395G^ enabled us to capture a structural state that escaped previous structural investigations, presumably due to its low abundance (Fig. 3d). This previously undescribed configuration reveals an inter-domain interaction network in which the conserved residue R465 in trans and the D1-D2 linker jointly coordinate communication with the Arg-finger R635 of the adjacent subunit through the ISS motif. This coupling modulates nucleotide coordination in the cis D2 active site. Beyond its direct catalytic effect, these findings highlight the central role of the D1-D2 linker in mediating inter-protomer communication. While previous biochemical analyses have implicated the linker in allosteric coupling between the two adjacent ATPase domains ^44,45^, our work reveals the detailed molecular basis of this communication network. A similar dependence on the linker has been described for the AAA+ ATPase Rix7, where deletion of its conserved N-terminal region causes severe growth defects ^46^, supporting inter-protomer communication via D1-D2 linker conserved across tandem AAA+ ATPases.

We further demonstrate that increased oxidative sensitivity of p97^D395G^ exacerbates its structural defects by enforcing a pronounced shift in the NTD conformation towards the ‘up’ state. Mass spectrometry uncovers two major sites for oxidative modification at key regulatory and catalytic loci: C184 at the NTD in the D1 interface and C522 within the D2 active-site pocket. Real-time NMR experiments on a p97^D395G^ construct lacking D2 confirm that oxidation at C184 alone is sufficient to drive this conformational shift. Given its position at the NTD-D1 interface, oxidative modification of Cys184 likely dysregulates NTD distribution, presumably influencing substrate processing by p97. Unexpectedly, all D2 nucleotide pockets in the oxidized p97^D395G^ maps lack nucleotide density despite the presence of ADP and P_i_ in D1 (Extended Fig. 9b). Our H_2_O_2_ treatment drove complete oxidation of C522, promoting its reorientation into the nucleotide-binding pocket, thereby blocking ATP binding and disrupting catalysis, in agreement with MDS data. Structurally, oxidized p97^D395G^ exhibits an upward shift of the D2 α_5_ helix, whereas oxidized p97^WT^ shows a downward displacement, indicating that α_5_ positioning regulates D2 catalytic activity by modulating nucleotide coordination through S652. These findings suggest that, within a dynamic cellular environment, reversible oxidative modification provides a tunable mechanism for controlling p97 activity.

Our study suggests that oxidative modification of C522 potentially provides a dynamic, redox-sensitive mechanism for tuning p97 ATPase activity. Reversible redox regulation of enzyme activity is well established in cellular systems ^47^. For example, oxidative modification of the catalytic cysteine transiently inhibits Caspase protease family ^48^. In this context, we propose that p97 function is actively regulated by redox signaling. Although C522 oxidation can be reversible under basal conditions, as demonstrated *in vitro* (Extended Fig. 7c), elevated ROS levels may promote irreversible oxidation at C522, potentially inactivating p97 and impairing protein homeostasis. Additionally, p97 negatively regulates the transcription factor, nuclear factor erythroid 2-related factor 2 (Nrf2), which is the master regulator of cellular response to oxidative stress, by extracting ubiquitinated Nrf2 from the Keap1-Cul3 E3 complex for proteasomal degradation ^49^. p97 suppression leads to increased levels of Nrf2 and its targeted gene products such as antioxidant response element genes ^50–52^. Oxidation of p97 at C522 inhibits its ATPase activity, which in turn may have an impact on Nrf2 levels. The mechanisms by which redox regulation of p97 differentially modulates Nrf2 signaling remains critical for understanding stress-response failure.

Our integrative strategy, combining NMR, MDS, MS, yeast genetics and cryo-EM provides an unprecedented view of how genetic mutation and oxidative stress converge into an unanticipated layer of control over p97 activity that is relevant under conditions of cellular stress and in neurodegenerative diseases. Collectively, these findings establish p97 as a redox-sensitive AAA+ ATPase whose activity is tuned by an evolutionarily conserved switch, thereby linking proteostasis to cellular stress responses.

## Methods

### Purification of p97 (full-length WT and mutants) and co-factor Ufd1-Npl4

GST-tagged full-length p97^WT^ was expressed from pGEX6p1-p97^WT^ construct, as it is described in ^15^. Briefly, cell lysate after sonication was incubated with Glutathione Sepharose beads (Cytiva) pre-equilibrated with PBS pH□7.4, 1□mM dithiothreitol (DTT). After washing, p97 was eluted with elution buffer (50□mM Tris-HCl, pH□8.0, 10 mM glutathione) and subjected to GST tag cleavage by HRV 3C protease. The protein was then applied to a Resource Q column (Cytiva) and eluted with a NaCl gradient (50□mM Tris-HCl, pH□8.0, 0–1□M NaCl), followed by further purification using a Superose 6 Increase column (Cytiva) in buffer containing 50 mM HEPES, pH 7.5, 150 mM NaCl, 5 mM MgCl_2_, and 0.5 mM Tris(2-carboxyethyl) phosphine (TCEP).

His-tagged p97^D395G^ mutant was expressed from pET28a-p97^D395G^ construct. Cell lysate was incubated with Ni^2+^-NTA beads pre-equilibrated with 50 mM Tris-HCl, pH 8.0, 150 mM NaCl. After washing with buffer containing 50 mM Tris-HCl, pH 8.0, 150 mM NaCl, p97^D395G^ was eluted with elution buffer containing 50 mM Tris-HCl, pH 8.0, 150 mM NaCl, 350 mM imidazole and subjected to His-tag cleavage by TEV protease. Following steps of ion-exchange chromatography (IEX) using Resource Q (Cytiva) and size-exclusion chromatography (SEC) were similar to the above-mentioned steps used for full-length p97^WT^.

All p97 mutations were introduced into the pGEX6p1-p97^WT^ construct using NEB site-directed mutagenesis kit (New England Biolabs), and mutant proteins were expressed and purified similarly to wild-type.

Ufd1 was expressed with a N-terminal His-tag using pET41b-His-Ufd1 obtained from Addgene ^53^, while untagged Npl4 was expressed using pET26b-Npl4 construct ^54^. To assemble the Ufd1-Npl4 (UN) complex, cell lysates of His-Ufd1 and untagged Npl4 were mixed, and the complex was purified using Ni^2+^-NTA beads. The lysate was incubated with Ni^2+^-NTA beads pre-equilibrated with 50 mM Tris-HCl, pH 8.0, and 100 mM NaCl. The beads were washed with 50 mM Tris-HCl, pH 8.0, 100 mM NaCl, and 25 mM imidazole. The heterodimer complex was eluted with elution buffer containing 50 mM Tris-HCl, pH 8.0, 100 mM NaCl, 300 mM imidazole, and 1 mM DTT. The protein was then applied to a Resource Q column (Cytiva) and eluted with a NaCl gradient (50□mM Tris-HCl, pH□8.0, 0–1□M NaCl), followed by further purification using a Superose 6 Increase column (Cytiva) in buffer containing 25 mM Tris-HCl, pH 8.0, 150 mM NaCl, 1 mM MgCl_2_, 5 mM β-mercaptoethanol (β-ME), and 10% glycerol.

### Purification of Ubiquitin and ubiquitinating enzymes E1 and E2-25K

Untagged human ubiquitin cloned into pGEX vector was expressed in *E. coli*. First, the cell lysate was applied to a pre-equilibrated DEAE (diethylaminoethyl) Sepharose column (Cytiva) (20 mM Tris-HCl, pH 8.0) to remove majority of proteins. Since ubiquitin has a relatively high isoelectric point (pI=9.0), it does not bind to the anion exchange column, and was therefore collected in the flowthrough. This was followed by pH adjustment using 1 M acetic acid to pH 4.0. The sample was then centrifuged at 20,000 rpm for 30 min at 4 °C to remove precipitations. The supernatant was collected and bound to pre-equilibrated SP (Sulphopropyl) Sepharose column (Cytiva) with equilibration buffer (20 mM acetate buffer, pH 4.0). After washing with equilibration buffer, ubiquitin was eluted with elution buffer containing 20 mM acetate buffer, pH 4.0, and 0.3 M NH_4_HCO_3._

Human E1 was expressed using pGEX6p1-E1-His8x construct as a dual fusion protein with N-terminal GST-tag and C-terminal His-tag. Firstly, the lysate was bound to Ni^2+-^NTA column pre-equilibrated with 50 mM Tris-HCl, pH 8.0, and 100 mM NaCl. After washing with washing buffer containing 50 mM Tris-HCl, pH 8.0, 100 mM NaCl, and 50 mM imidazole, E1 was eluted with elution buffer containing 50 mM Tris-HCl, pH 8.0, 100 mM NaCl, 300 mM imidazole, and 1 mM DTT. Following this, the elution fraction was bound to Glutathione Sepharose beads equilibrated with 50 mM Tris-HCl, pH 8.0, and 100 mM NaCl. After washing with buffer containing 50 mM Tris-HCl, pH 8.0, 100 mM NaCl, and 1 mM DTT, E1 was eluted with elution buffer containing 50 mM Tris-HCl, pH 8.0, 100 mM NaCl, 10 mM glutathione, and 1 mM DTT.

Human E2-25K was expressed using pET7-E2-25K-His construct with a N-terminal His-tag. It was bound to Ni^2+^-NTA column pre-equilibrated with 50 mM Tris-HCl, pH 8.0, and 100 mM NaCl. After washing with washing buffer containing 50 mM Tris-HCl, pH 8.0, 100 mM NaCl, and 50 mM imidazole, E2-25K was eluted with buffer containing 50 mM Tris-HCl, pH 8.0, 100 mM NaCl, 300 mM imidazole, and 1 mM DTT.

### Purification of model substrate Ub_4_-mEOS3.2

The Ub_4_-mEOS3.2 was purified as previously described ^55^. The protein contains a His-tag followed by a SUMO-tag on the N-terminal end and a His-tag on the C-terminal end. The lysate was bound to pre-equilibrated (50 mM Tris-HCl, pH 8.0, 150 mM NaCl) Ni^2+^-NTA column. After washing with buffer containing 50 mM Tris-HCl, pH 8.0, and 150 mM NaCl, His-SUMO-Ub_4_mEOS3.2-His fusion protein was eluted with elution buffer containing 50 mM Tris-HCl, pH 8.0, 150 mM NaCl, and 350 mM imidazole. Following this, Ulp1 protease was used for N-terminal SUMO-tag cleavage. The protein was then applied to a Resource Q column and eluted with a NaCl gradient (50□mM Tris-HCl, pH□8.0, 5mM β-ME, 0–1□M NaCl), followed by further purification using a Superose 6 Increase column (Cytiva) in buffer containing 25 mM Tris-HCl, pH 8.0, 150 mM NaCl, 1 mM MgCl_2_, and 0.5 mM TCEP.

### Ubiquitination reaction of p97 model substrate Ub_4_-mEOS3.2

Reactions were carried out at a final concentration of 40 μM Ub_4_-mEOS3.2, 0.2 μM E1, 10 μM E2-25K, 200 μM ubiquitin in 50 mM Tris-HCl, pH 8.0, 5 mM MgCl_2,_ 4 mM DTT and 5 mM ATP at 37 °C overnight. Ubiquitin and ATP were progressively added in small amounts every hour for a total duration of 6 hours. To purify ubiquitinated Ub_4_-mEOS3.2 fusion protein from the free ubiquitin chains, the reaction mixture was incubated with Ni^2+^-NTA resin and eluted with 300 mM imidazole and run over a Superose 6 column equilibrated with buffer containing 25 mM Tris pH 8.0, 150 mM NaCl, 1 mM MgCl_2_, 0.5 mM TCEP and 5% glycerol. Only the long chain ubiquitinated fractions were pooled together and concentrated with a centrifugal concentrator for further use in substrate unfolding assays.

### Photo-conversion of ubiquitinated p97 model substrate Ub_4_-mEOS3.2

The long chain ubiquitinated fractions were exposed to long wavelength UV (365 nm on a Vilber UV lamp, 230 V) for an hour while the sample was kept on ice. During the UV exposure step, the sample was checked for visible signs of precipitation and was briefly pipetted at regular intervals. The efficiency of photo-conversion was calculated based on the ratio between the absorbance at 508 nm/extinction coefficient at 508 nm and the absorbance at 573 nm/extinction coefficient at 573 nm.

### p97 substrate unfolding assay

Unfolding reactions were carried out following the protocol described in ^8^, in unfolding buffer (50 mM HEPES, pH 7.5, 100 mM NaCl, 10 mM MgCl_2_, 5 mM ATP, 1 mg/mL BSA) in presence of 20 nM ubiquitinated substrate, 500 nM p97 hexamer, and 500 nM UN. All components of the unfolding buffer including p97, UN and ubiquitinated substrate were preincubated in 384-well plate for 15 minutes at 37 °C before the addition of the ATP to initiate the reaction. Fluorescence was read on a TECAN SPARK plate reader using a 530 nm excitation filter (25 nm bandwidth) and 590 nm emission filter (25 nm bandwidth). Data was collected in replicates and was normalized to the initial reading.

### NMR spectroscopy

Perdeuterated and selectively I-δ_1_-[^13^CH_3_], V/L-γ_1_/δ_1_(*proR*)-[^13^CH_3_,^12^CD_3_] and M-ε_1_-[^13^CH_3_] labeled p97-ND1L for NMR samples were prepared as previously published ^15^. The buffer of protein samples for NMR spectroscopy was 25 mM HEPES, pH 7.5, 25 mM NaCl, and 5 mM TCEP in 100% D_2_O. Measurements were carried out at protein concentrations of 80-150 µM. Nucleotide states were prepared with 5 mM ADP, or 5 mM ATPγS (Jena Bioscience, Jena, Germany) with 4 mM MgCl_2_, respectively. Regeneration system measurements were prepared with 5 mM ATP, 4 mM MgCl_2_, 30–200 mM PEP, 4 mM ribose-5-phosphate, 50 mM KCl, and 10–50 U pyruvate kinase from *Bacillus stearothermophilus* (Merck, Darmstadt, Germany).

For oxidation, 12 mM H_2_O_2_ (Merck, Darmstadt, Germany) was added to the NMR sample of p97^D395G^ ND1L construct in the presence of 5 mM ADP and a time series of methyl-SOFAST HMQC spectra was acquired at 313 K.

2D methyl-TROSY spectra were recorded on a Bruker Avance III spectrometer operating at 800 MHz, employing a TCI cryoprobe. Experiments were performed at 310 K, 313 K, or 323 K, as indicated. Spectra were processed using TopSpin 3.7 and analyzed using CcpNmr v3 ^56^ or the Nmrglue suite ^57^. Chemical shift perturbations Δδ_methyI_ were determined as the Euklidean distance between the peak maxima in the 2D methyl-TROSY spectra according to 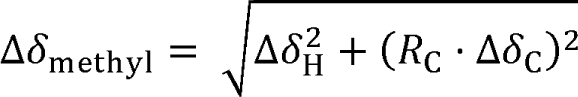. A scaling factor of R_C_ = 0.185 was used in the ^13^C dimension ^58^.

### Cryo-EM data acquisition

Quantifoil R2/1 Copper 200 micron or UltraAuFoil R2/2 Gold 200-micron grids were used for grid preparation. For ADP and ATPγS-bound datasets of p97^D395G^, purified p97^D395G^ was incubated in presence of ADP (#A2754 Sigma) and ATPγS (#NU-406 Jena Bioscience) for 15 minutes at room temperature. For p97^D395G^ in ADP.P_i_ state, p97 at 1.5□mg/ml was incubated in the ATP^reg^ system (4□mM ribose-5-phosphate, 4□mM MgCl_2_, 50□mM KCl, 13.3□U pyruvate kinase, 50□mM phospho-enol pyruvate, and 10□mM ATP) for 20□min at 37□°C. Octyl-beta-glucoside at a final concentration of 0.05% was added just before plunge-freezing. For oxidized p97 samples, protein was treated with 25 μM H_2_O_2_ and followed by SEC. Only hexameric fractions were used for making grids (Extended Fig. 7b). The oxidized samples were incubated in ATP^reg^ system before plunging. 3 μl of sample was pipetted onto the grids and then was rapidly plunge-frozen into liquid ethane/propane cooled by liquid nitrogen with a Vitrobot Mark IV (Thermo Fisher Scientific) in a chamber equilibrated at 10□°C with 100% humidity. Data acquisition was performed with Thermo Fischer Titan Krios G4 electron microscope at an acceleration voltage of 300 kV. Data sets were acquired using the program EPU software (V3.6) (Thermo Fischer Scientific) with Fringe-Free Imaging (FFI) and aberration-free image shift (AFIS) on a Falcon 4i Direct electron detector mounted after a Selectris energy filter with a slit width of 10 eV. Movies were acquired at x165,000 magnification (0.72 Å/pix).

### Cryo-EM image processing

For p97^D395G^ in ADP.P_i_ dataset, beam-induced motion correction was done using MotionCor2 ^59^ and CTF parameters were estimated using CTFFIND4 ^60^ both as implemented within Relion ^61^. The micrographs were subsequently imported into cryoSPARC ^62^ for further processing. A total of 368,013 particles were picked by crYOLO ^63^ and, 350,844 particles were subjected to heterogeneous refinement (C1 symmetry) after iterative rounds of 2D classification, resulting in two major classes: dodecameric (243,380 particles) and hexameric (124,663 particles). Owing to its higher resolution, the dodecameric particles were used for further processing. To improve local density, the dodecameric particles were computationally separated into individual hexamers without NTD, yielding 535,994 particles. Subsequent refinement resulted in a D1-D2 ring map at 2.92 Å. To further improve the NTD density, particles were subjected to 3D Flex refinement in cryoSPARC, resulting in an NTD-focussed reconstruction at 3.27 Å with improved density for the NTD. A composite map was generated by combining the D1-D2 ring-focused and NTD-focused maps. To analyze D1-D2 linker dynamics, focussed classification was carried out on C6 symmetry expanded particles of p97^D395G^ and p97^WT^ in ADP.P_i_ (EMD-16782) ^15^ dataset. A mask encompassing the D1 and D2 ATPase domain of one protomer and the D1 ATPase domain of the adjacent protomer in counter-clockwise direction was used (Fig. 3c, Supplementary video 3). This was followed by 3D classification which segregated two major classes in p97^D395G^ (Class I – 64% and Class II – 24%). In contrast, in p97WT four classes (Class i – 40%, Class ii– 6 %, Class iii – 19%, Class iv – 14%) were obtained but they were all similar to Class I of p97^D395G^. For all other data sets, motion correction and CTF estimation were performed in cryoSPARC. For the p97^D395G^ ADP dataset, 204,245 particles out of 282,981 initial particles were subjected to homogeneous refinement with C6 symmetry, yielding a 2.5 Å reconstruction. For the p97^D395G^ ATPγS dataset, 400,161 particles out of 524,112 initially picked particles were subjected to 3D classification with C1 symmetry, yielding classes with distinct NTD conformations. All classes displayed clear nucleotide density corresponding to ATPγS in the nucleotide binding pockets. Particles corresponding to NTD ‘up’ state (71,531 particles) were used for further refinement, resulting in a 2.2 Å reconstruction. For p97^WT^(Ox) in the ADP.P_i_ dataset, among 457,948 initial particles, 343,922 particles were subjected to 3D classification with C1 symmetry, yielding 2 major classes. These two classes were combined (204,214 particles) for further refinement using C6 symmetry, yielding a 2.46 Å reconstruction. For, p97^D395G^(Ox) ADP.P_i_ dataset, among 598,376 initial particles, 569,880 particles were subjected to 3D classification with C1 symmetry, yielding classes with distinct NTD conformations. Two classes were selected for an additional round of classification, resulting in 65,757 particles with all six NTDs in ‘down’ conformation. These particles were further refined with C6 symmetry, yielding a 2.81 Å reconstruction. C6 symmetry expansion was then performed, and focused classification and local refinement strategies around a single protomer improved the NTD density and revealed additional density in the D2 active site pocket consistent with oxidation of C522 as identified by MS. The remaining of the particles from 3D classification were selected and subjected for another round of 3D classification, enabling identification of different classes with varying NTD conformations such as 1 NTD down (69,273 particles), 2 NTD down (113,046 particles), 3 NTD down (50,566 particles), 4 NTD down (186,337 particles) and 5 NTD down (55,689 particles). Details of the cryo-EM experiments are provided in Supplementary Table 1-3.

### Model building

The published model 8OOI ^15^ was used as a starting point for initial model building of p97^D395G^ in ADP.P_i_ state, p97^D395G^ in ADP state, oxidized p97^WT^ in ADP.P_i_ state, and oxidized p97^D395G^ in ADP.P_i_. model^12^ was used a starting point for p97^D395G^ in ATPγS state. The model was first rigid-body docked into the map, followed by manual adjustment in Coot ^64^. The resulting model was refined using Phenix ^65^. Details of the model-building process and the quality of the final atomic models are provided in Supplementary Table 1-3.

### Surface plasmon resonance

For surface plasmon resonance measurement, biotinylated p97 protein was prepared with a C-terminal AviTag as described by ^66^. The binding affinities of nucleotides to p97 were measured on a Biacore X100 instrument. CM5 chips (Cytiva Life Sciences) were coated with NeutrAvidin by injecting a mixture of N-hydroxysuccinimide and 1-ethyl-3-(3-dimethylaminopropyl)-carbodiimide for 7 min, followed by a 7-min injection of 0.25 mg/mL NeutrAvidin. Finally, the surface was blocked by injecting ethanolamine (pH 8.3) for 2 min. Purified biotinylated p97-AviTag protein was immobilized in flow cell 2 to 1600 RU from HBS-P+ buffer (10 mM HEPES pH 7.4, 150 mM NaCl, 0.05% v/v surfactant P20) (Cytiva Life Sciences) with 5 mM MgCl_2_. 12-step 2-fold dilution series starting from 100 µM (ADP) or 50 µM (ATPγS) were prepared in HBS-P+ with 5 mM MgCl_2_, injected for 120 s at a flow rate of 30 µL/min, followed by regeneration with 500 mM NaCl in the same buffer. Response differences FC2-FC1 were fitted to a one-site kinetic model for ligand concentrations at which association and dissociation were sufficiently slow to resolve, allowing extraction of kinetic binding parameters.

### NADH-coupled ATPase assay

The ATPase rates of p97 were determined using an NADH-coupled ATPase assay where the oxidation of NADH is directly coupled to the rate of ATP hydrolysis and hence is used as a proxy to calculate the ATPase rate. Phosphoenolpyruvate (6□mM), NADH (1□mM), pyruvate kinase (1□U/100□μl), lactose dehydrogenase (1□U/100□μl) and purified protein (1–10 μM) were diluted into the ATPase buffer (25□mM HEPES, pH□7.5, 25□mM NaCl, 50□mM KCl, 4□mM MgCl_2_, and 0.5□mM TCEP) and distributed into a 96-well plate to a final volume of 120□μl. The reaction mixture was equilibrated at 37□°C for 10-15□min before the addition of 5 mM ATP. The rate of NADH consumption was monitored at 340 nm for 90 mins on a TECAN SPARK plate reader (TECAN). ATP-hydrolysis rates (ATP/min) were calculated based on at least three experimental replicates.

### Yeast genetic assays

All yeast strains were congenic with RTY1 ^67^ and generated by standard yeast genetic approaches. *HIS3-*marked plasmids encoding C-terminally V5-tagged yeast Cdc48 or the indicated point mutants were transformed into haploid yeast strains RTY863 (*cdc48*Δ*::natMX4*), RTY961 (*cdc48*Δ*::natMX4 otu1*Δ*::kanMX6*), RTY4187 (*cdc48*Δ*::natMX4 ufd1-2*), or RTY4193 (*cdc48*Δ*::natMX4 vms1*Δ*::kanMX6*) each harboring a copy of *CDC48* on a *URA3*-marked plasmid to maintain viability. Transformants were then struck on 5-fluoroorotic acid (5-FOA) media to evict the *URA3*-marked cover plasmid. To assess organismal health, 5-FOA evicts were then grown to mid-log in YPD media before back-dilution to OD_600_ = 0.1, and spotting in six-fold serial dilutions on the indicated media.

### *In-vitro* oxidation treatment using H_2_O_2_

All samples were oxidized with 25 μM or 50 μM H_2_O_2_ on ice for an hour. Following this, the samples were dialyzed overnight to remove traces of H_2_O_2_, followed by SEC to remove aggregated protein. The samples were then used for ATPase assays and for cryo-EM data collection.

### Mass spectrometry

To determine site-specific oxidative modifications via mass spectrometry, p97^WT^ and p97^D395G^ mutant treated with 25 µM H_2_O_2_ were used. An amount of 2.5 µg p97 samples were incubated in a total of 20µl 1x PBS buffer containing 1 mM EDTA and 100 mM N-ethylmaleimide to block cysteines. The reaction was carried out for 30 min at 22 °C and stopped by adding 1 µl 1 M Tris pH 7.5 and 5x sample buffer (62.5 mM Tris-HCl, pH 6.8, 0.35 M SDS, 50 % glycerol, bromophenole blue) without DTT followed by 10 % polyacrylamide gel separation. Protein-containing bands, which were stained with Coomassie brilliant blue, were cut out of the gel.

For in-gel digestion, gel pieces were destained by repeated addition of 50 mM ammonium bicarbonate and 50 % acetonitrile buffer with repeated swelling steps with 50 mM ammonium bicarbonate ^68^. The gel pieces were then shrunk with 500 µl pure acetonitrile. Samples were not reduced with DTT and directly alkylated with 55 mM iodoacetamide in 100 mM ammonium bicarbonate for 20 min at 22 °C in the dark. Supernatant was removed and gel pieces were shrunk with 500 µl pure acetonitrile. Tryptic digestion was carried out for 16 hours at 37 °C and 350 rpm in 10 mM ammonium bicarbonate with 10 % acetonitrile and 13 ng/µl trypsin. The peptides were extracted by adding 5 % formic acid in acetonitrile (2 times the volume of the digestion), and the supernatant was transferred to a new tube. A second extraction step with the same volume as the digestion was performed on the gel pieces. Both supernatants were combined and dried in a SpeedVac (Eppendorf® Concentrator Plus). For desalting, samples were dissolved in 50 µL of 0.1 % trifluoroacetic acid, loaded onto C18 Spin Tip, washed twice with 20 µL of 0.1 % trifluoroacetic acid and eluted with 2x 20 µL of 0.1 % trifluoroacetic acid in 80 % acetonitrile.

Peptide samples were analyzed on an Orbitrap Fusion Tribrid mass spectrometer (Thermo Scientific) and interfaced with an Easy-nLC 1200 nanoflow liquid chromatography system (Thermo Scientific). The samples were reconstituted in 0.1 % formic acid in 5 % acetonitrile and loaded onto the C18 analytical column (50 μm × 15 cm, Thermo Scientific). Peptides were dissolved at a flow rate of 300 nL/min using a linear gradient of 6-44% solvent B (0.1% formic acid in 80% acetonitrile) for 45 minutes. Data-dependent acquisition with full scans over a range of 320-1850 m/z was performed with the Orbitrap mass analyzer at a mass resolution of 60,000 in the normal mass range, with custom-defined target value for automatic gain control and an automatic injection time. Only peptides with charge states 2-10 were used and the dynamic exclusion was set to 60 seconds. Precursor ions were fragmented using higher-energy collision dissociation (HCD) with normalized collision energy (NCE) set to 30 %. An intensity threshold of 5.0e4 with an isolation window of 1.6 m/z was defined. The fragment ion spectra were recorded with a resolution of 30,000, the target value for the automatic gain control was set to standard and the injection time mode was dynamic. Each sample was measured in technical duplicates.

Raw files of LC-MS/MS measurements were analyzed with MaxQuant (version 2.1.4.0) ^69^ with standard parameters if not stated otherwise. The following variable modifications were considered: acetylation (N-terminus), oxidation (methionine), carbamidomethylation (cysteine), glutathionylation (cysteine), di-oxidation (cysteine), tri-oxidation (cysteine), NEM (cysteine), NEM+water (cysteine) and Asp→Gly. The minimum peptide length was set to 4, the maximum peptide length was set to 35 and the alignment between runs as well as the label-free quantification (LFQ) were enabled. For more specific analysis, p97 amino acid sequence was used as search template and the match-between-runs option was set. The ratio of peptide intensities (peptide variant with respective cysteine oxidation / all other cysteine modifications) was used to determine the relative intensity of the respective modification in the peptides.

## Molecular Dynamics Simulations

### System model

The reduced p97^WT^ structure was built from cryo-EM coordinates in the up state derived from PDB entry 7LMY ^18^. The engineered mutations A232E and E578Q present in the deposited structure were reverted to the WT sequence. ATP and one magnesium ion were placed in each of the twelve nucleotide binding sites. Missing residues were modeled using the python Modeller plugin ^70^. The system topology was generated with psfgen ^71^. The structure was solvated in a 0.15 M NaCl solution to neutralize the total charge within a cubic simulation box of 28×28×13 nm^3^. Energy minimization and equilibration were performed for 200 ns under NPT conditions. Reduced p97^D395G^ was generated from the equilibrated WT structure, whereas oxidized p97^WT^ was generated from the unequilibrated WT structure by mutating residues D395G and C522 in each subunit, respectively. Afterwards, systems were resolvated and ionized with 0.15 M NaCl.In oxidized p97, the C522 residue was mutated to cysteine-sulfonic acid using parameters available in the additive CHARMM36 force field for non-standard amino acids ^72^ . Structural data from all three systems were collected every 0.1 ns during a 1 μs NPT production simulation. Trajectories were visualized and structures rendered in VMD 1.9.4 ^71^.

### Molecular dynamics simulation

The solvated protein systems were simulated using NAMD version 3.0b3 ^73^ with proteins and ions described by the CHARMM36m force field ^74^ and water molecules represented by the TIP3P model ^75^. Periodic boundary conditions were employed throughout the simulations. Atomic trajectories were propagated using a verlet integration scheme with a time step of 2 fs. Nonbonded interactions were computed using van der Waals and short-range electrostatic potentials for atom pairs separated by less than 1.2 nm, with a switching function applied between 1.0 and 1.2 nm. Long-range electrostatic interactions were calculated on a grid with 0.12 nm spacing using the PME method ^76^. Interaction pair lists were constructed for atom pairs within 1.4 nm and updated every 200 fs. Covalent bonds involving hydrogen atoms were constrained using the SHAKE algorithm ^77^, whereas water geometry was maintained with the SETTLE algorithm ^78^. Temperature regulation was achieved using a Langevin thermostat ^79^ with a damping coefficient of 1 p/s, maintaining the system at 310 K. Pressure was controlled isotropically at 1 atm using a Nosé-Hoover Langevin piston ^80^ with an oscillation period of 200 fs and a damping time of 100 fs.

### Contact number analysis

Contacts within p97 were quantified using contact numbers ^81^ derived from the minimum periodic distances *d* between heavy atoms *k* and *l*.

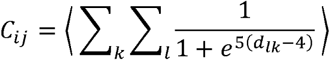

Angle brackets denote the time average over the trajectory frames. Interatomic distances below 0.4 nm□yield contact numbers between 1 and 0.5, whereas larger separations result in values approaching zero. Due to the homo-hexameric structure of p97, contact numbers calculated from all six subunits were averaged to enhance statistical sampling.

### C522 distance to active center

A geometric center was defined using the C_α_ coordinates of residues 520, 521, 524, 525, 526, 543, 577, 578, 685, and 688, together with residues 609, 632, 635, 636, 637, and 638 from the neighboring subunit. For each trajectory frame, the distance between the sulfur atom of C522 and this geometric center was calculated.

### Oxidation likelihood

Principal component analysis was conducted using the cartesian coordinates of the C_α_ atoms of residues surrounding C522 together with five atoms from the nearby ATP molecule for the reduced p97^WT^ and reduced p97^D395G^ trajectories. Energetic minima were identified from the conformational distributions projected along the first and second principal components. Seven representative conformations located within and adjacent to these minima were selected, and the free energy change associated with oxidation of C522 to deprotonated cysteine sulfonic acid was calculated using thermodynamic integration. Due to the large size of the fully solvated hexamer, calculations were performed on a reduced subsystem consisting of two neighboring monomers A and F along with their bound ATP molecules. To maintain the structural integrity of the complex during the alchemical transformation, all C_α_ atoms were harmonically restrained to their minimized coordinates using a force constant of 5 kcal/mol/Å^2^. Because the oxidation introduced a net change in charge, a sodium ion positioned 30 Å away from the protein was simultaneously transformed from a neutral to a charged particle to preserve overall charge neutrality. The thermodynamic integration protocol and simulation parameters were adopted from Mauck *et al* ^82^.

Analysis of the oxidation free energies, revealed two distinct groups separated by approximately 20 kcal/mol. However, projection onto the principal component-based collective variables failed to separate these groups within the conformational ensemble. To overcome this limitation, a custom collective variable was defined as a linear combination of the cartesian coordinates of the same atom set used for the principal component analysis (Extended Fig. 9d). The coefficients of this linear combination were optimized using the minimize function implemented in SciPy ^83^ to maximize the separation between oxidation favorable and oxidation unfavorable conformations while minimizing variance within each group:

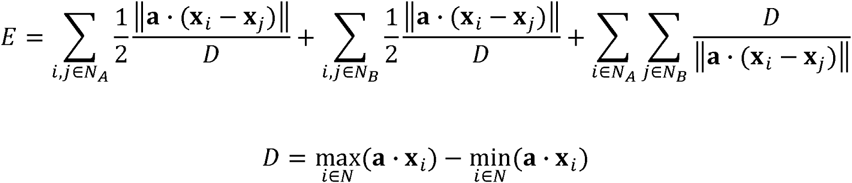

Here, *E* denotes the error function, **a** the collective variable, **x** the feature vector, and *N_A_*and *N_B_* indicate the two groups of the oxidation free energy values. An optimized collective variable was obtained that distinguished between the two conformational groups. As an independent validation, the oxidation free energy starting directly from the oxidized cryo-EM structure was determined to be energetically between the two groups. Projection of this structure onto the optimized collective variable was in agreement with the alchemical free energy results and positioned between the two conformational groups. The optimized collective variable was subsequently applied to all frames from the p97^WT^ and p97^D395G^ trajectories, enabling the determination of probability density distributions that quantify the relative populations of oxidation-favorable conformational states.

## Acknowledgements

This work was supported by the German Research Foundation (DFG) through SFB1690 (Project A02 to E.S.), Germany’s Excellence Strategy (EXC 2067/1–390729940 to E.S.), the Emmy Noether program (Project 394455587 to A.K.S.), and the Rise Up! program of the Boehringer Ingelheim Foundation (BIS) to A.K.S. Additional support was provided by the German Research council (grants 496470458, 516836828, and 550938463 to F.S.) and by the NIH (R01GM118600 to R.J.T., Jr.). The support was also provided by JSPS/MEXT KAKENHI (JP18H05501 to SF). The authors gratefully acknowledge the scientific support and HPC resources provided by the Erlangen National High Performance Computing Center (NHR@FAU) of the Friedrich-Alexander-Universität Erlangen-Nürnberg (FAU) under the NHR project b235bb_2. NHR funding is provided by federal and Bavarian state authorities. NHR@FAU hardware is partially funded by the German Research Foundation (DFG) – 440719683. Cryo-EM instrumentation was funded by the DFG Major Research Instrumentation program and the Ministry of Science and Culture of the State of Lower Saxony (448415290 to E.S.).

## Author contribution

Protein samples were produced by SRR, MD, and MS. SRR, MD, MS and YS performed and analyzed the biological and biophysical experiments. SRR and TCC performed data acquisition and data processing for cryo-EM experiments. MD and MS performed NMR experiments and analysis. MK and TAM conducted MD simulation. MW and SRR performed the MS experiments. MD and GW performed the SPR experiment. SA and AAN performed yeast experiments. MK, MZ, RTJ, FS, SF, AKS and ES designed and supervised the research. Figures were prepared by SRR, MD, MW, and MK. SRR, MD, MK, AKS, and ES wrote the manuscript with contributions from all authors.

