## Supplementary Figure for "The vacuolar tauopathy-associated mutation D395G confers redox sensitivity to p97/VCP"

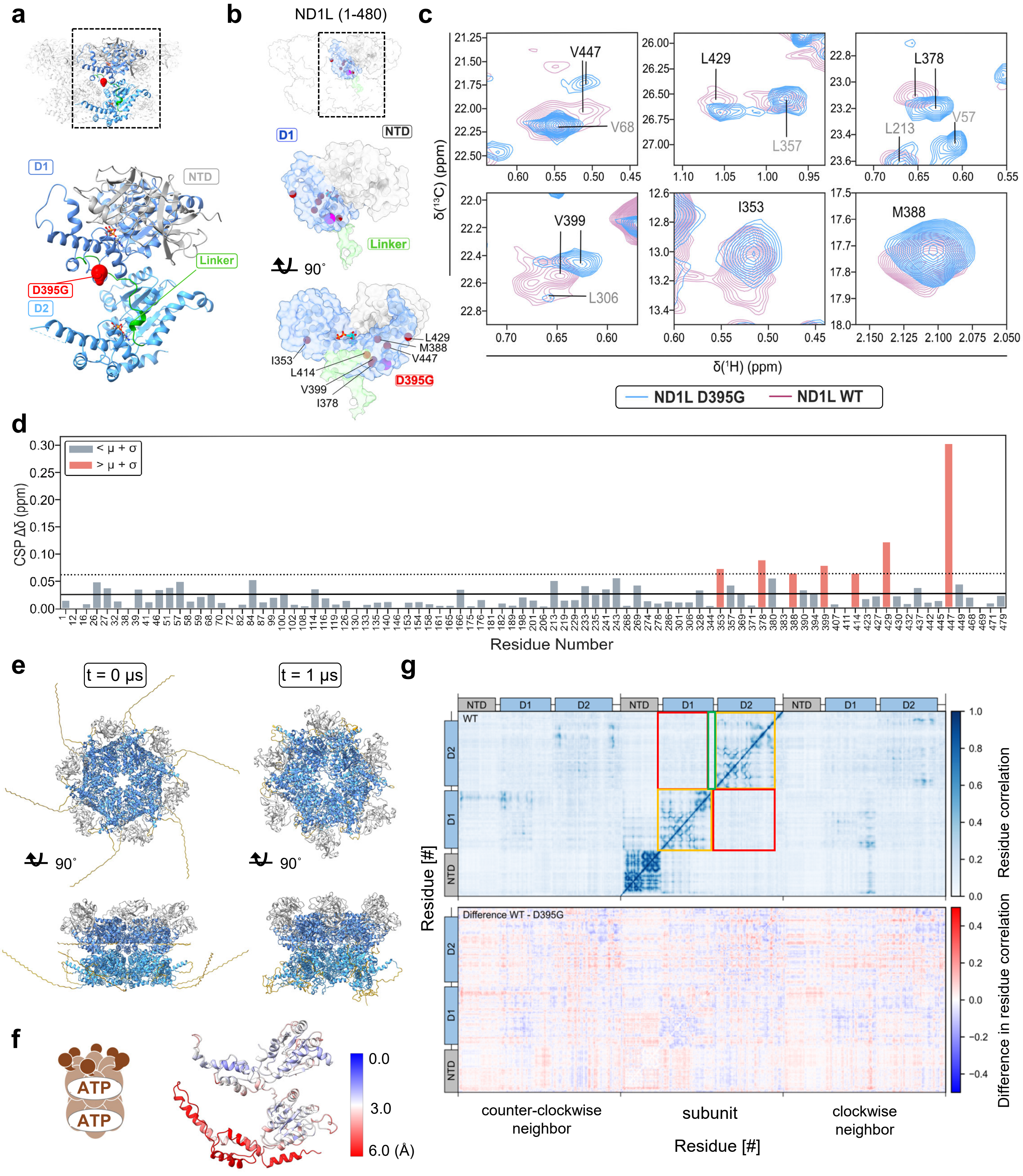

a
D1
NTD
Linker
D395G
D2
b
ND1L (1-480)
D1
NTD
Linker
90˚
L429
M388
V447
I353
L414
V399
I378
D395G
c
V447
V68
L429
L357
L378
L213
V57
V399
L306
I353
M388
δ(¹³C) (ppm)
δ(¹H) (ppm)
ND1L D395G
ND1L WT
d
< μ + σ
> μ + σ
CSP Δδ (ppm)
Residue Number
e
t = 0 μs
t = 1 μs
90˚
90˚
f
ATP
ATP
0.0
3.0
6.0 (Å)
g
NTD
D1
D2
WT
Residue correlation
Difference WT - D395G
Difference in residue correlation
Residue [#]
counter-clockwise neighbor
subunit
clockwise neighbor
Residue [#]

**Extended Data Figure 1: Structural comparisons of p97$^{WT}$ and p97$^{D395G}$.**

**a,** Ribbon representation of full-length p97 with the D395G mutation highlighted as red sphere; NTD (grey), D1 (cyan), D1–D2 linker (green), and D2 (blue). Nucleotides are shown in ball-and-sphere representation. **b,** p97$_{1\text{-}480}$ ND1L structure used for NMR experiments highlighting all available methyl probes as spheres. D395G mutation site is shown as a large red sphere and methyl groups displaying CSPs (chemical shift perturbations) are highlighted in pink. **c,** Overlay of selected regions from methyl HMQC spectra of p97 ND1L$^{WT}$ and ND1L$^{D395G}$, both in the ADP state, highlighting the signals that display the largest CSP effects from D395G mutation. **d,** Quantitative analysis of CSPs between p97$^{D395G}$ and p97$^{WT}$ in the ADP state. Only those residues are shown for which methyl groups could be unambiguously assigned to NMR signals. Highlighted in red are CSPs above the threshold for significance, defined as the mean CSP plus one standard deviation ($\mu + \sigma$); $\mu$= 0.0088 ppm, $\sigma$ = 0.0076 ppm, $\mu + \sigma$ = 0.0164 ppm. Euclidean distance calculations and weighting factors are described in the Methods section. **e,** *Left:* The full-length p97$^{WT}$ (PDB:7LMY) was used as the starting structure for MDS. Missing sequences (shown in golden yellow) were built with Modeller. *Right:* The equilibrated p97$^{WT}$ structure after 1 μs of MDS. **f,** Backbone RMSD between trajectory-averaged MD structures of p97$^{WT}$ and p97$^{D395G}$ protomers with ATP bound in D1 and D2 domains. **g,** Correlation maps for p97$^{WT}$ and the difference between p97$^{WT}$and p97$^{D395G}$. Correlations between neighboring-subunits are shown relative to the central subunit, with values ranging from fully correlated (1) to uncorrelated (0). Intradomain correlations within the D1 and D2 (orange) are observed, with no interdomain correlations (red), while the D1-D2 linker region shows correlation specifically with the D2 domain (green). In the difference map, red and blue regions denote stronger correlations in p97$^{WT}$ and p97$^{D395G}$, respectively.

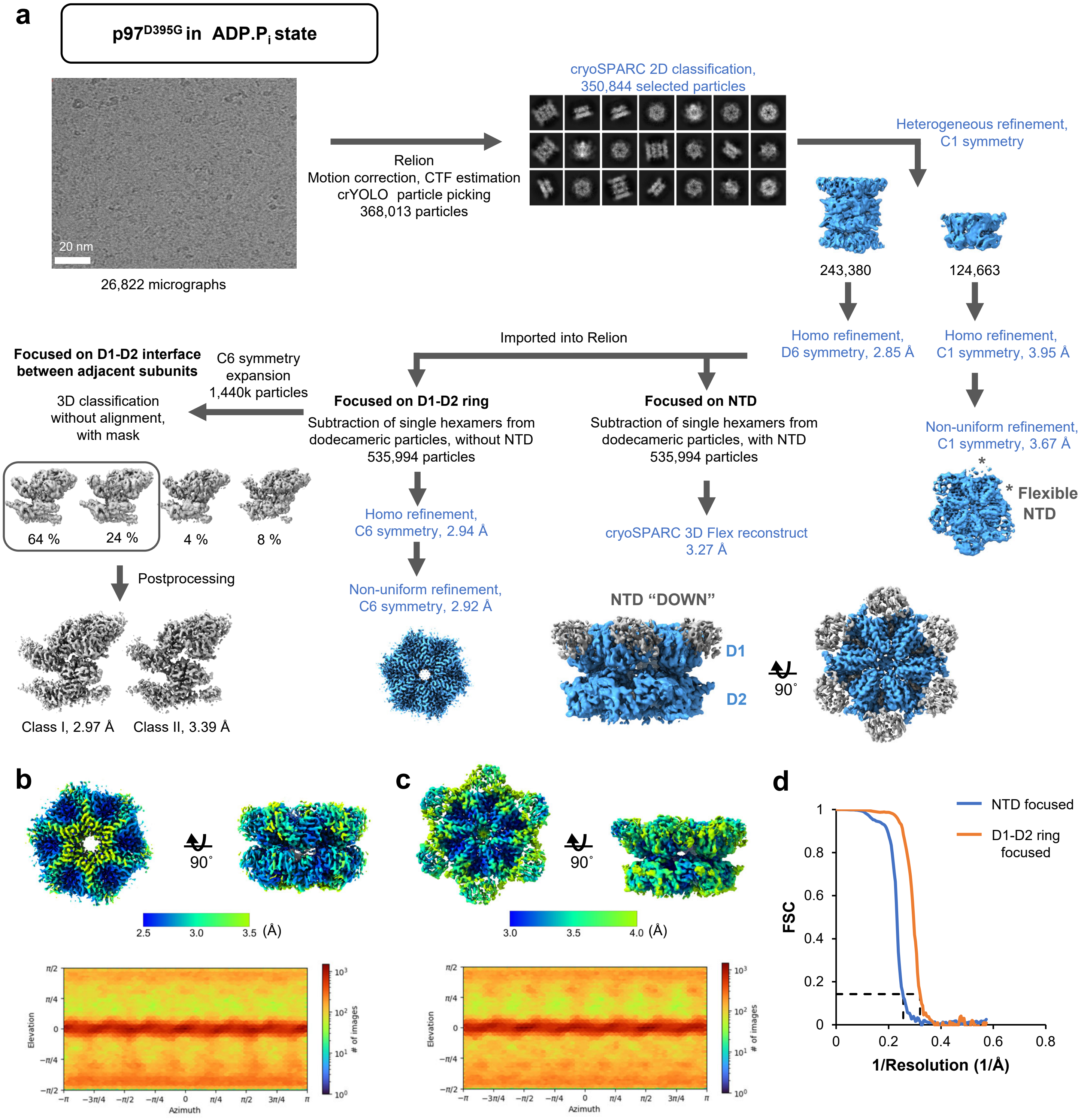


**Extended Data Figure 2: Data processing workflow for p97$^{D395G}$ in ADP.P$_i$ state**

**a,** Classification of p97$^{D395G}$ in an ATP$^{reg}$ system (ADP.P$_i$ state) yielded both hexameric and dodecameric assemblies. Owing to its higher resolution, the dodecameric class was used for further processing. To improve the resolution, dodecameric particles were computationally separated into constituent hexamers, followed by focused classification and refinement of the D1-D2 ring and the NTD. Both of the focused maps were combined to generate a composite map. Subtracted particles without the NTD were used for focused classification with a mask encompassing D1 and D2 ATPase domains of one protomer and D1 ATPase domain of adjacent protomer in counter-clockwise direction. 3D classification without alignment yielded 2 major classes, which were further processed to generate a Class I map at 2.97 Å and Class II at 3.39 Å. The same approach was applied to previously published data for p97$^{WT}$, as described in Extended Fig. 6 Processing steps performed in RELION are indicated in black, where as those carried out in cryoSPARC are in blue. **b, c** Local resolutions of the D1-D2 ring-focused reconstruction (b) and the NTD-focused reconstruction (c) are displayed in gradation according to the indicated scale. The corresponding particle orientation distribution is shown below. **d,** Resolutions were determined on the basis of gold standard Fourier Shell Correlation between independently refined half maps (FSC = 0.143, dashed line).

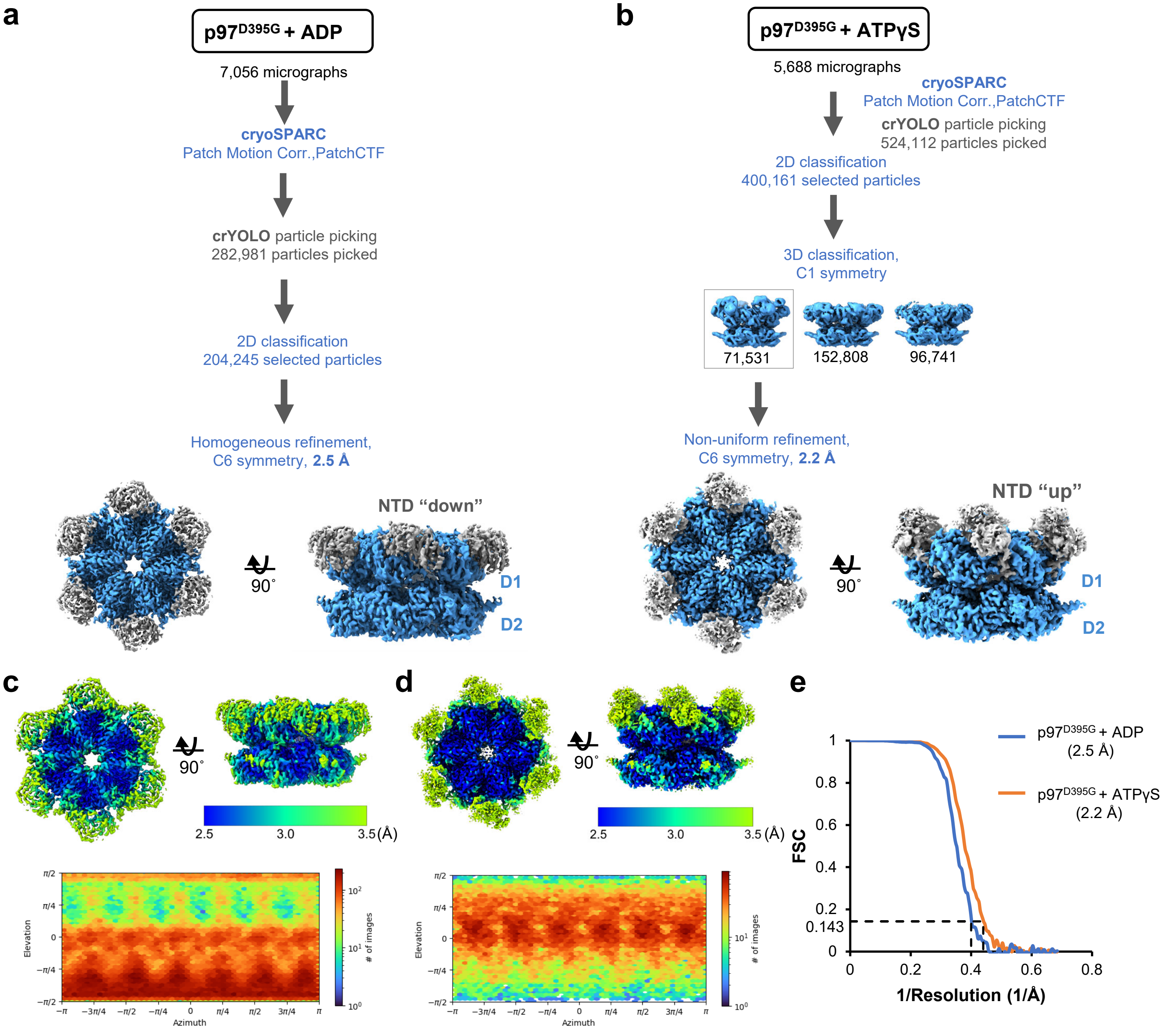


**Extended Data Figure 3: Data processing workflow for p97$^{D395G}$ in presence of ADP and ATPγS**

**a,** Flowchart showing classification and refinement procedures used for obtaining the reconstruction of ADP-bound p97$^{D395G}$. **b,** Flowchart showing classification and refinement procedures used for obtaining the reconstruction of ATPγS-bound p97$^{D395G}$. Upon 3D classification with C1 symmetry, particles with NTDs in different positions were obtained. NTD conformations other than 'up' state have been seen for p97$^{WT}$ in the presence of ATPγS when processed with C1 symmetry. Since all classes here showed density corresponding to ATPγS in the nucleotide binding pocket, hence the particles with NTD 'up' state were only used for further refinement. **c, d** Local resolutions of ADP-bound p97$^{D395G}$ (c) and ATPγS-bound p97$^{D395G}$ (d) are displayed in gradient according to the indicated scale. The corresponding particle orientation distribution is shown below. **e,** Resolutions were determined on the basis of gold standard Fourier Shell Correlation between independently refined half maps (FSC = 0.143, dashed line).

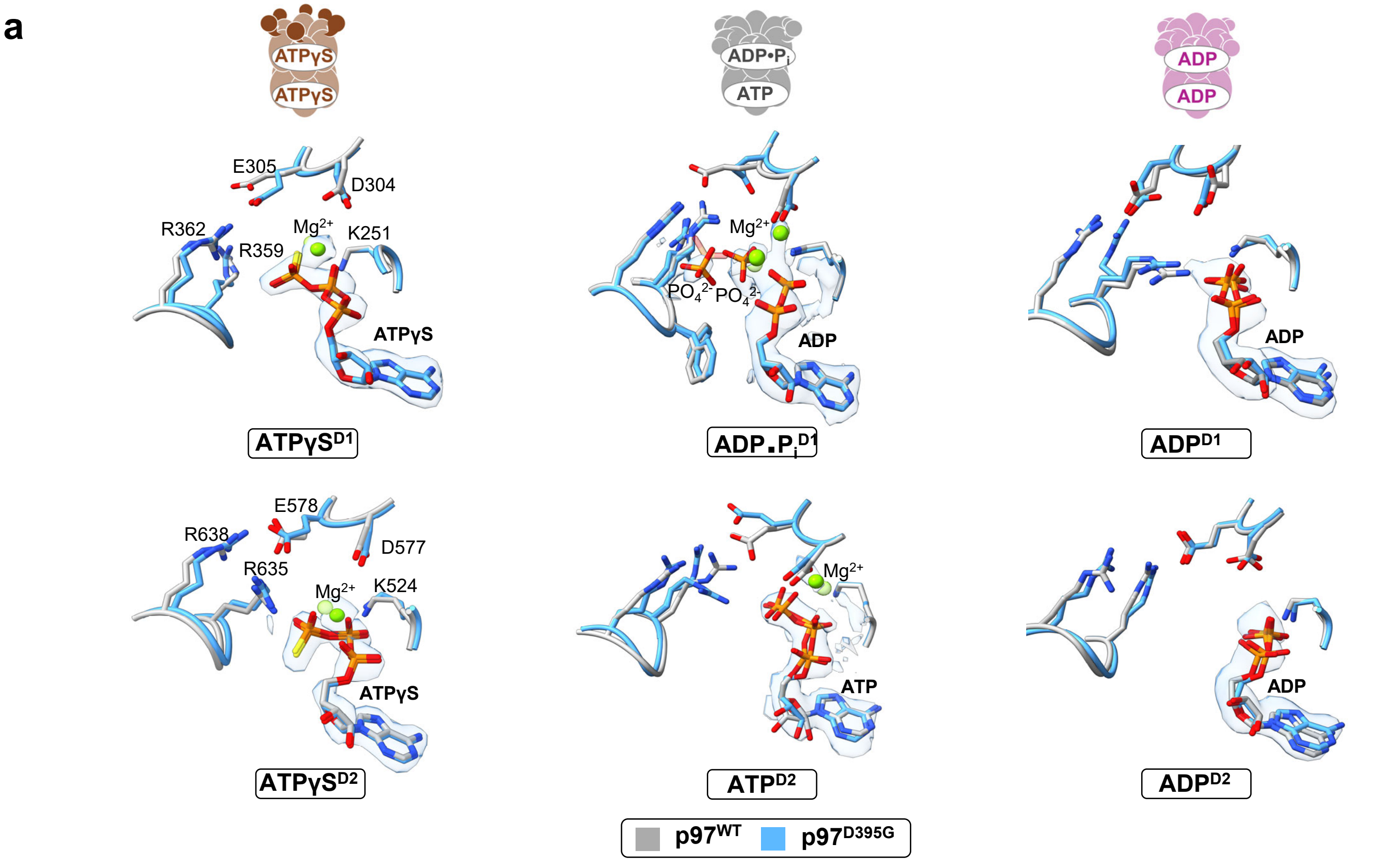


**Extended Data Figure 4: Active site pockets of p97$^{D395G}$ in ADP.Pi$^{D1}$ state shows heterogeneity**

**a,** D1 and D2 ATPase active-site pockets of p97$^{D395G}$ in different nucleotide-bound states, shown together with the corresponding cryo-EM density for the bound nucleotide. p97$^{D395G}$ (blue) in the ATPγS-bound, ADP-bound, and ADP·P$_i$-bound states is superimposed on the reported p97$^{WT}$ models in the ATPγS-bound (PDB: 5FTN), ADP-bound (PDB: 5FTK), and ADP·P$_i$-bound (PDB: 8OOI) states. The magnesium and phosphate ions for p97$^{WT}$ structures are rendered transparent for distinction.

**a**

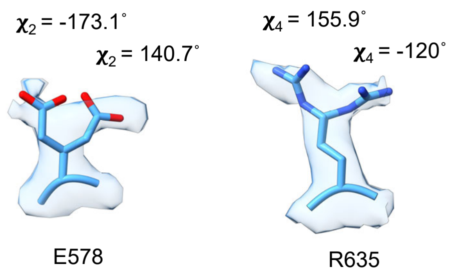


**b**

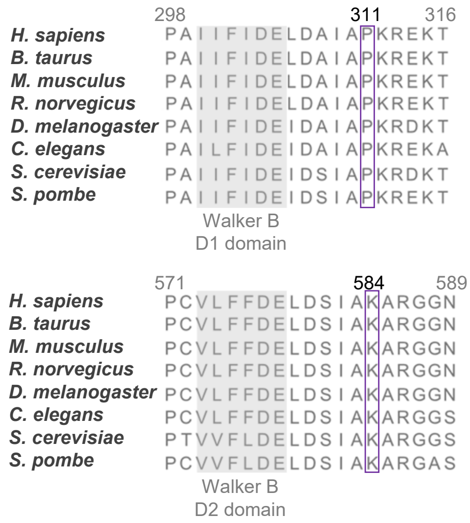


**c**

| p97 | Nucleotide | $k_a$ (x $10^5$ $M^{-1}s^{-1}$) | $k_d$ ($s^{-1}$) | $K_D$ (nM) |
|---|---|---|---|---|
| WT | ADP | 3.2–3.8 | 0.1–0.2 | 339–533 |
| | ATPγS | 3.0–4.7 | 0.05–0.08 | 174–182 |
| D395G | ADP | 4.3–4.5 | 0.12–0.16 | 288–361 |
| | ATPγS | 4.2–4.3 | 0.03–0.04 | 66.2–86.9 |

**d**

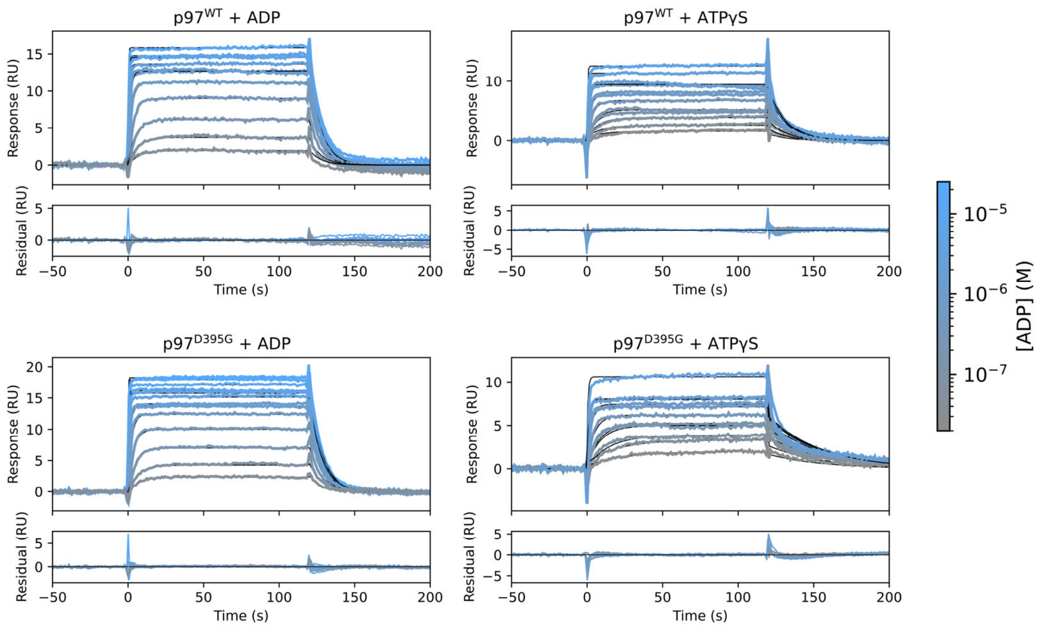


**Extended data Figure 5: D395G mutation affects D2 active site but not D1**

**a,** Double side chain rotamers of D578 and R635 in the D2 domain of p97$^{D395G}$ shown with cryo-EM density map. **b,** Multiple sequence alignment showing K584 in p97 is unique to the D2 ATPase domain, while P311 exists in the equivalent position in D1 ATPase domain. **c,** Kinetic binding parameters of p97$^{WT}$ and p97$^{D395G}$ for ADP and ATPγS determined in two technical replicates using surface plasmon resonance (SPR). **d,** Representative SPR sensorgrams (response difference FC2–1) of nucleotide binding immobilized p97, used to determine kinetic binding parameters in panel C. Fit curves to a kinetic one-site binding model are shown in black, residuals display the difference between sensorgrams and fits. Binding contributions from the D1 and D2 domains could not be resolved. Sensorgrams for the following nucleotide concentrations are displayed and were used in the analysis: p97$^{WT}$ + ADP: 9 step 2-fold dilution series from 12.5 µM to 0.049 µM; p97$^{WT}$ + ATPγS: 9 step 2-fold dilution series from 6.25 µM to 0.0244 µM with duplicates of 0.195 µM and 0.391 µM; p97$^{D395G}$ + ADP: 10 step 2-fold dilution series from 25 µM to 0.049 µM; p97$^{D395G}$ + ATPγS: 8 step 2-fold dilution series from 3.125 µM to 0.0244 µM with duplicates of all concentrations except for the three highest. Experiments performed in two technical replicates yielded affinities of the same order of magnitude for p97$^{WT}$ and p97$^{D395G}$, indicating that the reduced catalytic activity is not due to altered nucleotide affinity.

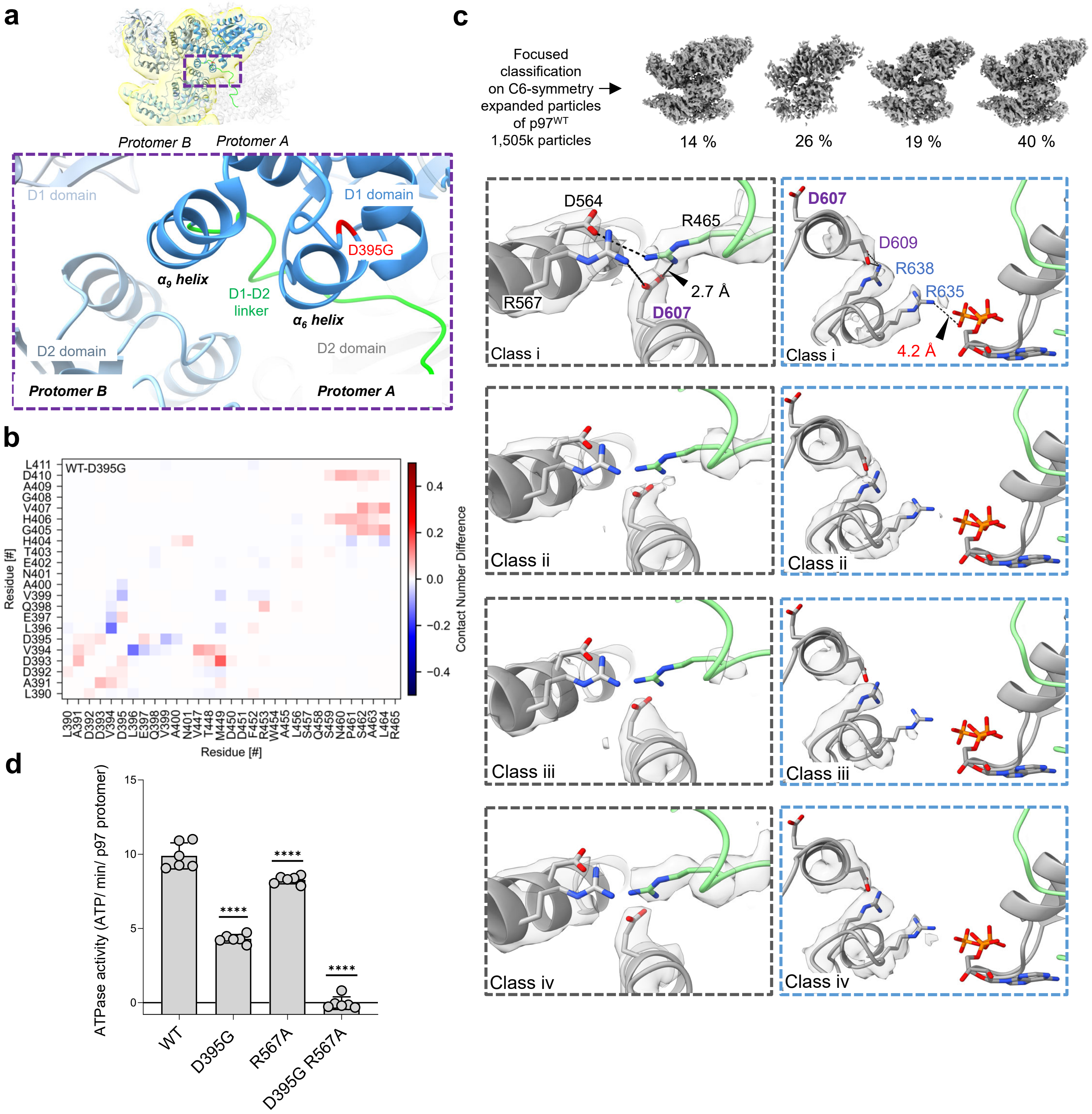


**Extended Data Figure 6: Structural basis of allosteric communication via D1-D2 linker**

**a,** The location of $\alpha_6$ helix harboring the D395G mutation (red), along with the adjacent $\alpha_9$ helix, which extends as D1-D2 linker (lime) on its C-terminal end. The mask used for focused classification is shown in yellow. The inset shows a zoomed-in view of the interaction interface between the D1-D2 linker of protomer A (blue) and the D2 domain of protomer B (cyan). **b,** Contact differences from MDS between $p97^{WT}$ and $p97^{D395G}$ for residues surrounding the extended D395G region. Red and blue colors indicate contacts more frequent in $p97^{WT}$ and $p97^{D395G}$, respectively. **c,** Focused classification, using a mask as described for $p97^{D395G}$, on published $p97^{WT}$ data yielded 4 classes similar to the Class I of $p97^{D395G}$, characterized by R465-D607 and D609-R638 interaction and thereby weakening interaction of R635 with nucleotide. The cryo-EM density maps corresponding to key residues of all classes of $p97^{WT}$ is shown here at the same contour threshold. **d,** ATPase assays shows alanine substitution of a conserved D1-D2 interface residue R567 modestly reduces ATPase activity, possibly by altering the interactions at the D1-D2 interface. However, upon introduction of R567A mutation in the background of D395G mutation, activity is completely abolished demonstrating a coupled effect of the two mutations on overall p97 activity. All measurements include at least 3 technical replicates (Asterisks indicate statistical significance calculated using one-way ANOVA: $P < 0.0001$).

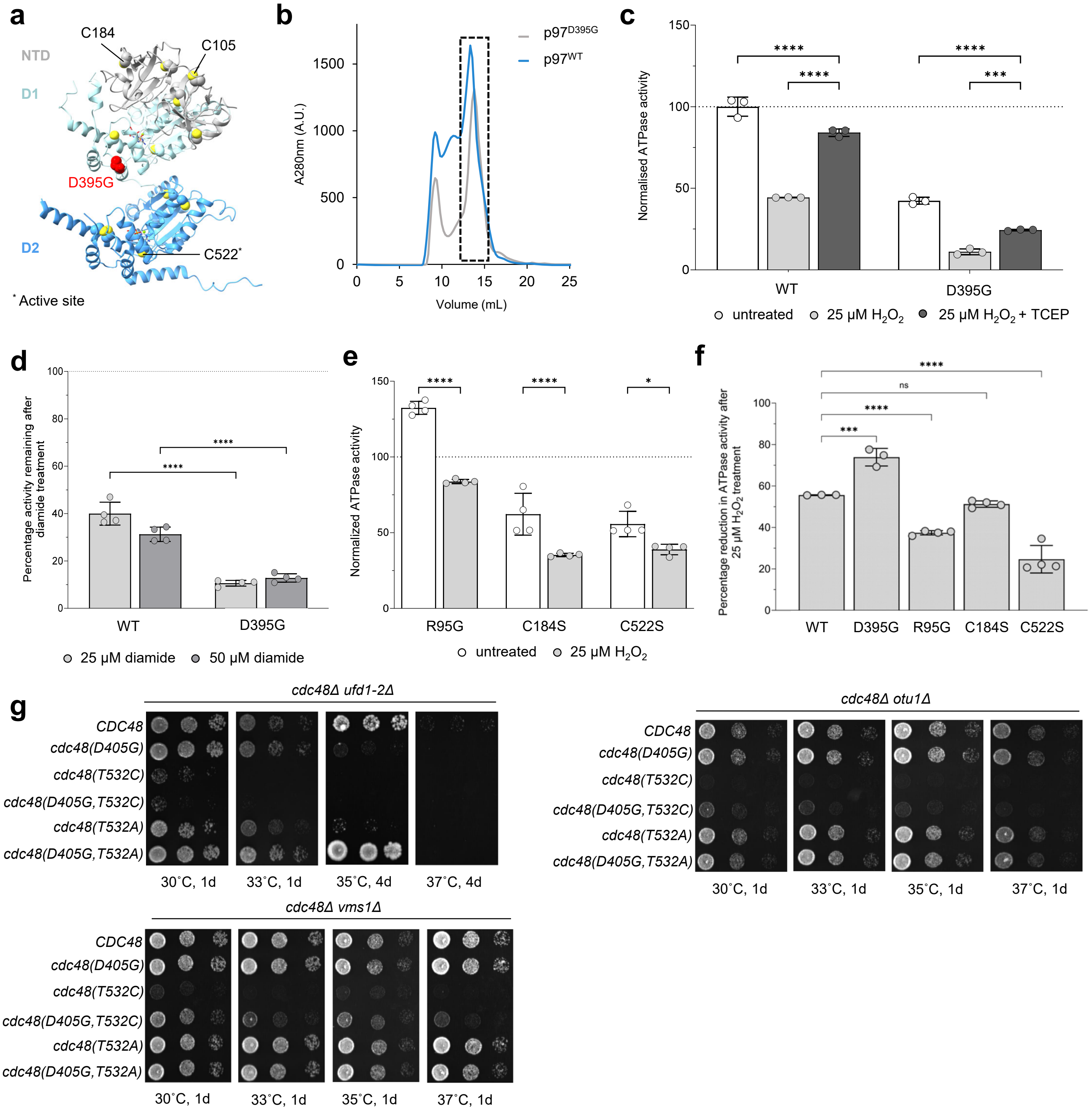


**Extended Data Figure 7: D395G mutation enhances sensitivity to oxidation**

**a,** The positions of the twelve cysteine residues within a single p97 protomer are shown as yellow spheres and D395G mutation as red sphere. **b,** Size exclusion chromatograms of p97 hexamers after oxidation treatment with 25 µM $H_2O_2$. Hexameric p97 fractions (dashed box) were utilized for subsequent ATPase assays and cryo-EM data acquisition. **c,** After treatment with 25 µM $H_2O_2$, the recovery of ATPase activity in $p97^{WT}$ and $p97^{D395G}$ was measured following addition of the reducing agent TCEP. $p97^{D395G}$ exhibited reduced recovery compared with $p97^{WT}$, with all values normalized to the activity of untreated $p97^{WT}$. **d,** ATPase activity remaining after diamide treatment relative to $p97^{WT}$ untreated. The dashed line denotes untreated $p97^{WT}$ activity. **e,** ATPase activity of p97 mutant proteins after treatment with 25 µM $H_2O_2$ normalized to untreated $p97^{WT}$ activity. **f,** Reduction in ATPase activity of oxidized p97 mutants after 25 µM $H_2O_2$ treatment, shown as a percentage decrease relative to the untreated condition. $p97^{D395G}$ exhibits the greatest reduction compared to MSP-1 mutant $p97^{R95G}$, highlighting differential susceptibility. Cysteine-to-serine substitution mutants generated based on MS data, show that following 25 µM $H_2O_2$ treatment C522S displays the minimum reduction in ATPase activity indicating that it confers resistance to oxidation. Substitution with serine preserves the D2 ATPase activity, which is the major contributor to overall p97 activity. All measurements include at least 3 technical replicates (Asterisks indicate statistical significance calculated using two-way ANOVA: ns, not significant, $P < 0.05$ (*), $P < 0.001$(***), $P < 0.0001$(****)). **g,** Yeast growth assays at elevated temperatures reveal genetic interactions between the Cdc48 T532C, T532A, and D405G substitutions and mutations in Cdc48 cofactors, including Ufd1, Vms1, and Otu1.

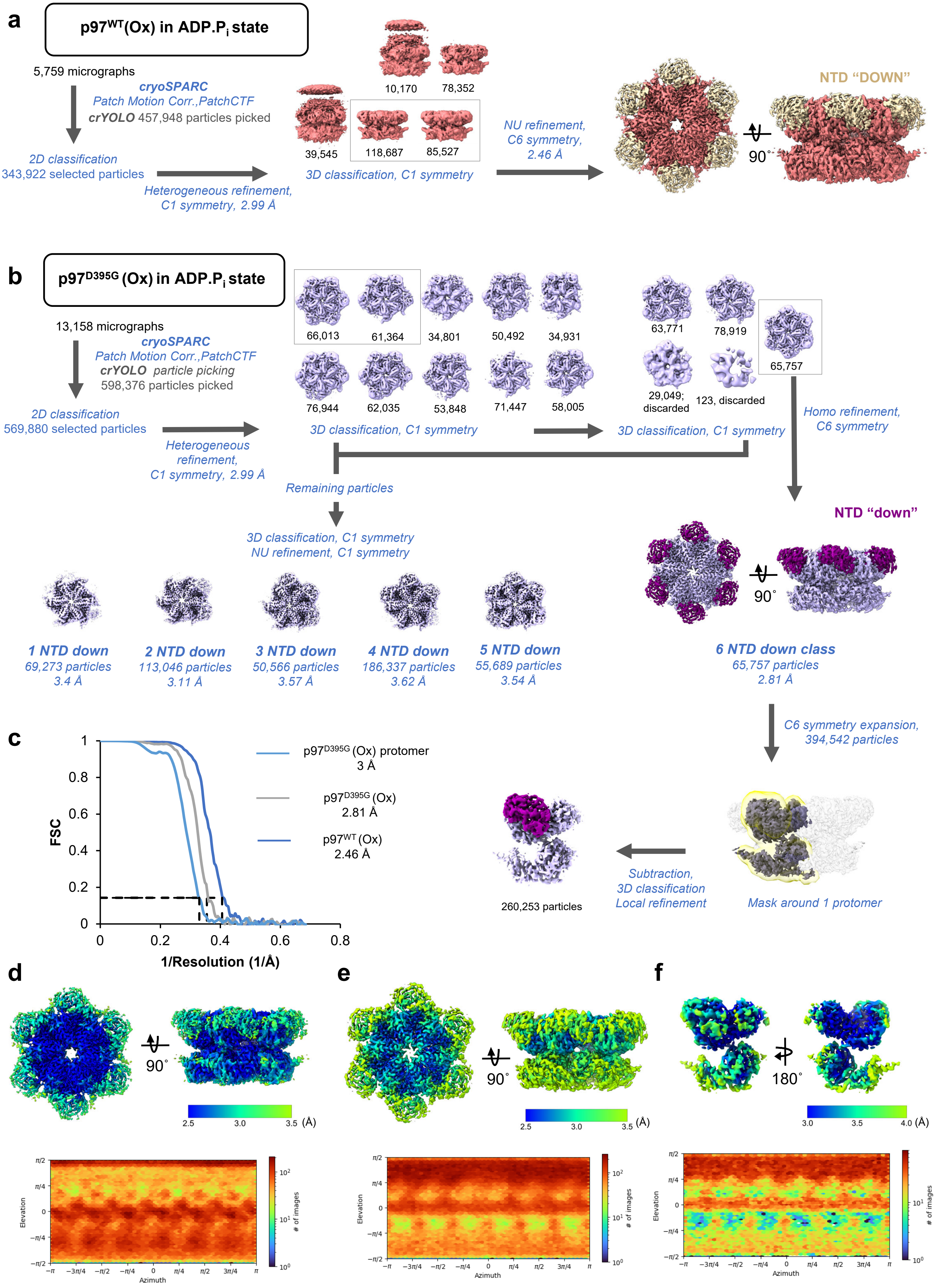
a
p97WT(Ox) in ADP.Pi state
5,759 micrographs
cryoSPARC
Patch Motion Corr.,PatchCTF
crYOLO 457,948 particles picked
2D classification
343,922 selected particles
Heterogeneous refinement,
C1 symmetry, 2.99 Å
10,170
78,352
39,545
118,687
85,527
3D classification, C1 symmetry
NU refinement,
C6 symmetry,
2.46 Å
NTD "DOWN"
90˚
b
p97D395G (Ox) in ADP.Pi state
13,158 micrographs
cryoSPARC
Patch Motion Corr.,PatchCTF
crYOLO particle picking
598,376 particles picked
2D classification
569,880 selected particles
Heterogeneous refinement,
C1 symmetry, 2.99 Å
66,013
61,364
34,801
50,492
34,931
76,944
62,035
53,848
71,447
58,005
3D classification, C1 symmetry
63,771
78,919
65,757
29,049; discarded
123, discarded
3D classification, C1 symmetry
Homo refinement,
C6 symmetry
Remaining particles
3D classification, C1 symmetry
NU refinement, C1 symmetry
NTD "down"
90˚
1 NTD down
69,273 particles
3.4 Å
2 NTD down
113,046 particles
3.11 Å
3 NTD down
50,566 particles
3.57 Å
4 NTD down
186,337 particles
3.62 Å
5 NTD down
55,689 particles
3.54 Å
6 NTD down class
65,757 particles
2.81 Å
C6 symmetry expansion,
394,542 particles
c
p97D395G (Ox) protomer
3 Å
p97D395G (Ox)
2.81 Å
p97WT (Ox)
2.46 Å
FSC
1/Resolution (1/Å)
260,253 particles
Subtraction,
3D classification
Local refinement
Mask around 1 protomer
d
90˚
2.5
3.0
3.5 (Å)
e
90˚
2.5
3.0
3.5 (Å)
f
180˚
3.0
3.5
4.0 (Å)
Elevation
Azimuth
### of images

**Extended Data Figure 8: Data processing workflow for p97$^{WT}$ (Ox) and p97$^{D395G}$ (Ox) in ATP$^{reg}$ system**

**a,** Flowchart showing classification and refinement procedures used for obtaining the reconstruction of oxidized p97$^{WT}$ in presence of ATP$^{reg}$. **b,** Flowchart showing classification and refinement procedures used for obtaining the reconstruction of oxidized p97$^{D395G}$ in presence of ATP$^{reg}$. Multiple rounds of classification and refinements were carried out with C1 symmetry to separate out the different classes of particles on the basis of NTD position. Particles with all NTDs in 'down' position were further refined with C6 symmetry to improve the density. C6 symmetry-expanded particles from the all-NTD 'down' class were subjected to local refinement with a mask encompassing a single protomer to further improve the NTD density. Subsequent 3D classification resolved a class exhibiting additional density consistent with oxidation adjacent to C522. **c,** Resolutions were determined on the basis of gold standard Fourier Shell Correlation between independently refined half maps (FSC = 0.143, dashed line). **d-f,** Local resolutions of p97$^{WT}$ (Ox) in ATP$^{reg}$ system (d), all-NTD 'down' class of p97$^{D395G}$ (Ox) in ATP$^{reg}$ (e), and p97$^{D395G}$ (Ox) from all-NTD 'down' class (f) are displayed in gradient according to the indicated scale. The corresponding particle orientation distribution is shown below.

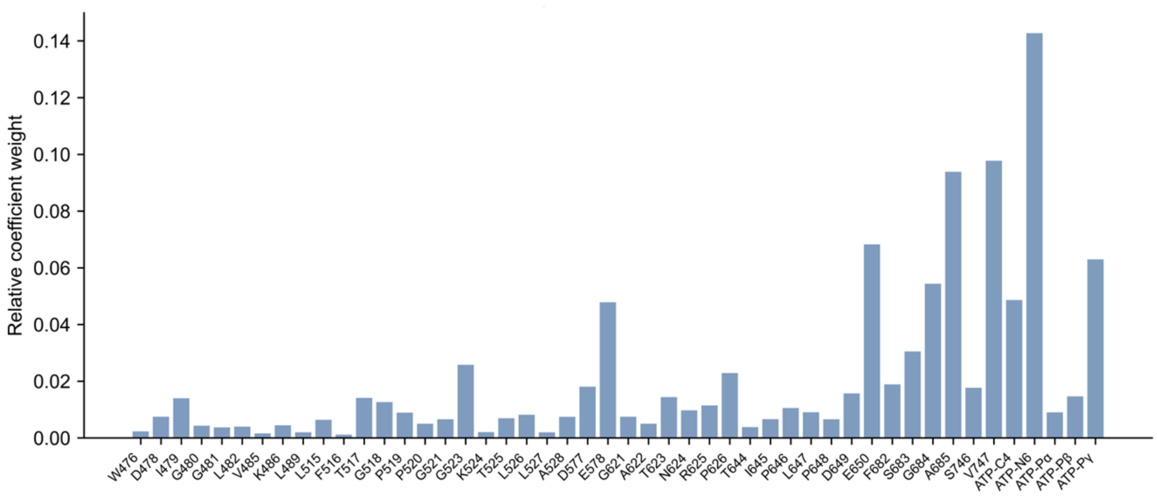


**Extended Data Figure 9: Weight coefficients in the oxidation collective variable**
Normalized relative atomic weight coefficients for the optimized collective variable described by coordinates of residue $C_\alpha$ atoms and selected ATP atoms. Cartesian coordinate coefficients for each atom are summed and represented as single values. Fluctuations or rearrangements of residues with higher coefficients correlate more strongly with changes in the oxidation free energy of C522.

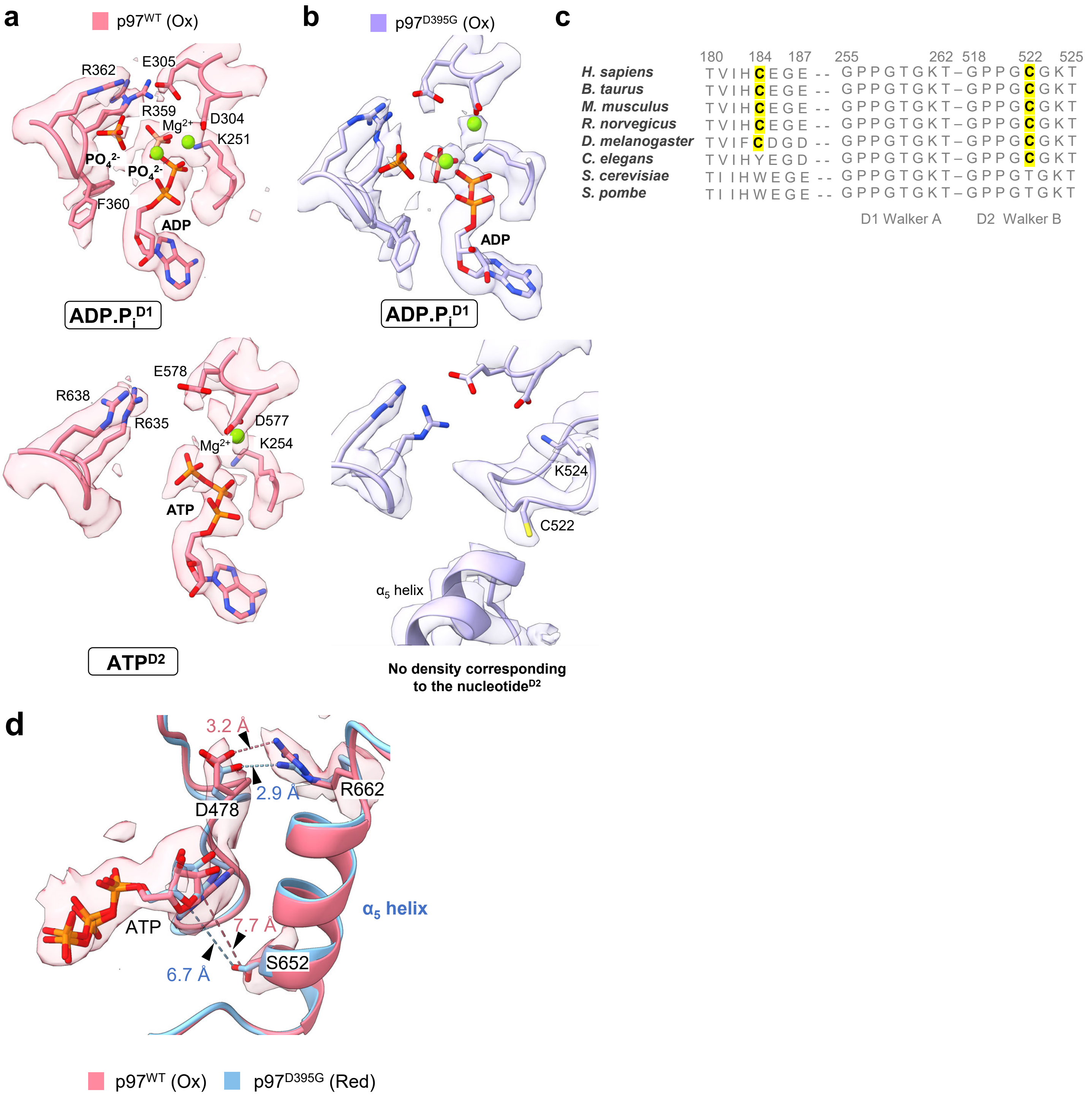


**Extended Data Figure 10: p97$^{WT}$ (Ox) shows $\alpha_5$ helix shift reminiscent of p97$^{D395G}$ (Red)**

**a,** D1 and D2 ATPase nucleotide-binding pockets of p97$^{WT}$ (Ox), shown with the corresponding cryo-EM density. **b,** D1 and D2 ATPase active-site pockets of p97$^{D395G}$ (Ox), shown with the corresponding cryo-EM density. **c,** Multiple sequence alignment showing conservation of C522 across higher eukaryotes, indicating the evolutionary acquisition of a cysteine residue in the catalytic D2 pocket. **d,** D2 active site configuration in p97WT(Ox) resembles that of p97D395G (Red), exhibiting improper nucleotide coordination by S652 due to a downward α5 helix shift.
